# Genetic Consequences of Social Stratification in Great Britain

**DOI:** 10.1101/457515

**Authors:** Abdel Abdellaoui, David Hugh-Jones, Kathryn E. Kemper, Yan Holtz, Michel G. Nivard, Laura Veul, Loic Yengo, Brendan P. Zietsch, Timothy M. Frayling, Naomi Wray, Jian Yang, Karin J.H. Verweij, Peter M. Visscher

## Abstract

Human DNA varies across geographic regions, with most variation observed so far reflecting distant ancestry differences. Here, we investigate the geographic clustering of genetic variants that influence complex traits and disease risk in a sample of ~450,000 individuals from Great Britain. Out of 30 traits analyzed, 16 show significant geographic clustering at the genetic level after controlling for ancestry, likely reflecting recent migration driven by socio-economic status (SES). Alleles associated with educational attainment (EA) show most clustering, with EA-decreasing alleles clustering in lower SES areas such as coal mining areas. Individuals that leave coal mining areas carry more EA-increasing alleles on average than the rest of Great Britain. In addition, we leveraged the geographic clustering of complex trait variation to further disentangle regional differences in socio-economic and cultural outcomes through genome-wide association studies on publicly available regional measures, namely coal mining, religiousness, 1970/2015 general election outcomes, and Brexit referendum results.

## Introduction

The first law of geography states that “everything is related to everything else, but near things are more related than distant things”.^1^ Humans living near each other tend to share more ancestry with each other than with humans that live further away, which is reflected in genome-wide patterns of genetic variation on a global scale^2^ and on finer scales.^3–5^ Regional differences in allele frequencies are driven by genetic drift (i.e., the random fluctuations of allele frequency each generation), natural selection pressures, migrations, or admixture (i.e., two previously isolated populations interbreeding). Out of these four mechanisms, genetic drift is the only mechanism not expected to disproportionately affect genetic variants that are associated with heritable human traits. Natural selection targets heritable traits over extended periods of time, thereby affecting allele frequencies of the genetic variants that are associated with the traits under selection. Earlier studies have identified natural selection pressures on many trait-associated variants by looking for extreme allele frequency differences between different ancestries.^3,6,7^ Migration is behavior, and since most behavioral traits have heritable components,^8^ migration is likely to be associated with genetic variants that influence behavior. Long-distance migratory events may in turn result in admixture. Internal migrations (i.e., migrations within countries) may lead to geographic clustering of trait-associated genetic variants beyond the clustering of ancestry and may occur for a variety of reasons. They may be driven by the search for specific neighborhood, housing, and inhabitant characteristics, and/or socio-economic factors (e.g., education or job-related considerations),^9^ such as the mass migrations from rural to industrial areas during the industrialization.^10^ These geographic movements may coincide with regional clustering of heritable social outcomes such as socio-economic status and major group ideologies (e.g., religion^11^ and political preference^12^).

Understanding what drives the geographic distribution of genome-wide complex trait variation is important for a variety of reasons. Studying regional differences of genetic variants associated with complex traits that reflect education, wealth, growth, health, and disease, may help understand why those traits are unevenly distributed across Great Britain. Besides the known regional differences in income and SES, significant regional differences have been reported for mental^13^ and physical^14^ health problems. Regional differences in wealth and health are likely linked to each other,^15–17^ and have been shown to be partly driven by migration.^14,18^ If genome-wide complex trait variation is geographically clustered, this should also be taken into account in certain genetically-informative study designs. Mendelian randomization for example uses genetic variants as instrumental variables to identify causality, under the assumption that the genetic instrument is not associated with confounders that influence the two traits under investigation.^19^ Geographic clustering of genetic complex trait variation could introduce gene-environment correlations that violate this assumption.^20^ Such gene-environment correlations could also introduce bias in heritability estimates in twin and family studies,^21^ and could affect signals from genome-wide association studies (GWASs). Furthermore, studying the genetics of migration and geographically clustered cultural phenomena that are related to how society is organized, such as SES, political preference, and religiosity, may help us to further understand regional differences beyond what can be learned from standard observational data. For example, as we will show in this study using a novel regional GWAS approach, we can compute genetic correlations between these clustered social phenomena and a wide range of other traits for which GWASs have been conducted through their GWAS summary statistics.^22^ This can teach us about how these regional differences are related to traits that have not been measured in the same dataset.

In this study, we first investigate whether genome-wide complex trait variation is geographically clustered after accounting for ancestry differences; if so, this may reflect the genetic consequences of more recent (internal) migration events. In addition, we investigate whether genome-wide complex trait variation is sufficiently clustered to capture the heritability of regional cultural outcomes such as coal mining, religiousness, and political preference by conducting GWASs on publicly available regional measures. We will then utilize the genetic signals from these GWASs to estimate genetic correlations between the regional measures and a wide range of complex traits.

## Data and Analysis

We investigated the geographic clustering of ancestry and complex trait variation using genome-wide single-nucleotide polymorphism (SNP) data from ~450,000 British individuals of European ancestry from the UK Biobank project.^23^ Ancestry within Great Britain was captured by conducting a principal component analysis (PCA)^24^ on genome-wide SNPs, a method that has been shown to successfully capture ancestry differences within relatively homogeneous populations.^3^ Genome-wide complex trait variation was captured by polygenic scores, which are created by weighting an individual’s alleles by the estimated allelic effects on the trait of interest and then summing the weights, resulting in predictive scores for each individual. We built polygenic scores for 456,426 individuals from 1,312,100 autosomal SNPs using effect estimates from 30 published GWASs on traits related to psychiatric disease, substance use, personality, body composition, cardiovascular disease, diabetes, reproduction, and educational attainment (see Supplementary Table 1). Importantly, the 30 GWASs that produced the effect estimates did not include UK Biobank participants.^25^ Geographic clustering of genetic variation was then investigated using 320,940 unrelated individuals and their birthplace by testing whether the spatial autocorrelation (Moran’s *I*) is significantly greater than zero for ancestry-informative principal components (PCs), polygenic scores, and the residuals of polygenic scores after regressing out the first 100 PCs. The spatial autocorrelation (Moran’s *I*) is the correlation in a measure among nearby locations in space, and its values range between −1 (dispersed) to 0 (spatially random) to 1 (spatially clustered).^26^ Supplementary Figure 1 shows geographic locations of UK Biobank participants. Furthermore, we test whether polygenic scores that showed significant geographic clustering were associated with an index of economic deprivation of the neighborhood (the Townsend index) and migration into or out of the most economically deprived regions (coal mining areas), while accounting for ancestry differences (100 PCs).

We subsequently investigate whether geographic clustering of genome-wide complex trait variation is associated with regional cultural outcomes by running genome-wide association analyses on coal mining, regional estimates of the proportion of religious vs non-religious inhabitants, election outcomes of the Brexit referendum and of the 1970 & 2015 general elections. We estimate the degree to which these regional differences share genetic influences with a range of traits related to cognitive ability, socio-economic status (SES), personality, behavior, substance use, mental and physical health, well-being, reproduction, and body composition.

For more detailed descriptions of the data and analyses, see Online Methods.

## Geographic Clustering of Genome-Wide Ancestry and Complex Trait Variation

In line with earlier studies,^5^ British ancestry showed significant geographic clustering: the first 100 genetic PCs all show Moran’s *I* statistics that are greater than 0, with 72 PCs showing an empirical *p*-value < .0005, the Bonferroni corrected threshold, and 95 PCs showing an empirical *p*-value < .05. Many PCs roughly capture the differentiation between Scotland, England, and Wales (see Figure 1 for the first 5 PCs; see https://holtzyan.shinyapps.io/UKB_geo/ for maps of all 100 PCs). The geographic distributions of the ancestry differences captured by the PCs are likely to reflect consequences of historical demographic events.^5^ These include old population movements and settlements, followed by generations of relatively isolated (sub)populations that went through genome-wide allele frequency differentiation through genetic drift and, perhaps, differential natural selection pressures.

**Figure 1:**
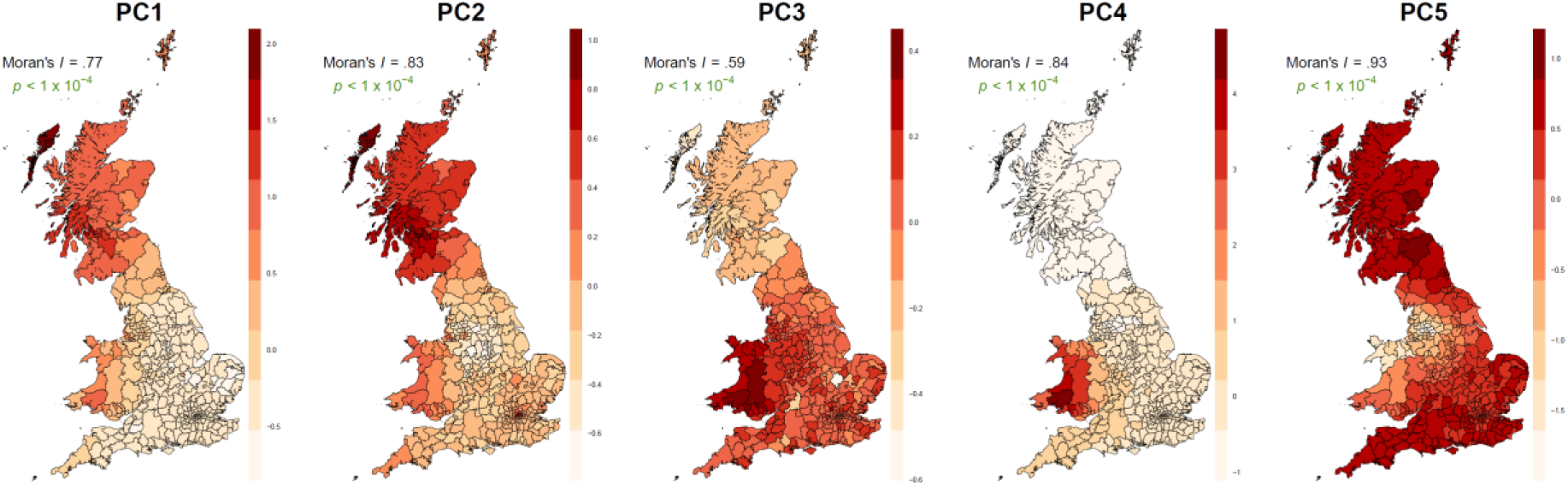
The geographic distributions (birthplace) of the first five PCs, Moran’s *I* and empirical *p*-values for Moran’s *I*. *P*-values denoted in green are significant after Bonferroni correction.

Without controlling for ancestry, 27 out of the 30 polygenic scores tested showed a Moran’s *I* significantly greater than 0, indicating significant geographic clustering (Figures 2 & 3, see https://holtzyan.shinyapps.io/UKB_geo/ for maps of all polygenic scores). Only age at menarche, agreeableness, and caffeine consumption were not significantly geographically clustered. Many clustered polygenic scores showed geographic distributions that were similar to the ancestry differences captured by the PCs. After regressing out the 100 ancestry-informative PCs, 16 polygenic scores remained significantly geographically clustered with FDR correction, with educational attainment (EA) showing the highest Moran’s *I* (before PC correction: Moran’s *I* = .57, empirical *p* < 10^−4^; after PC correction: Moran’s *I* = .51, empirical *p* < 10^−4^; see Figures 3 & 4).

**Figure 2:**
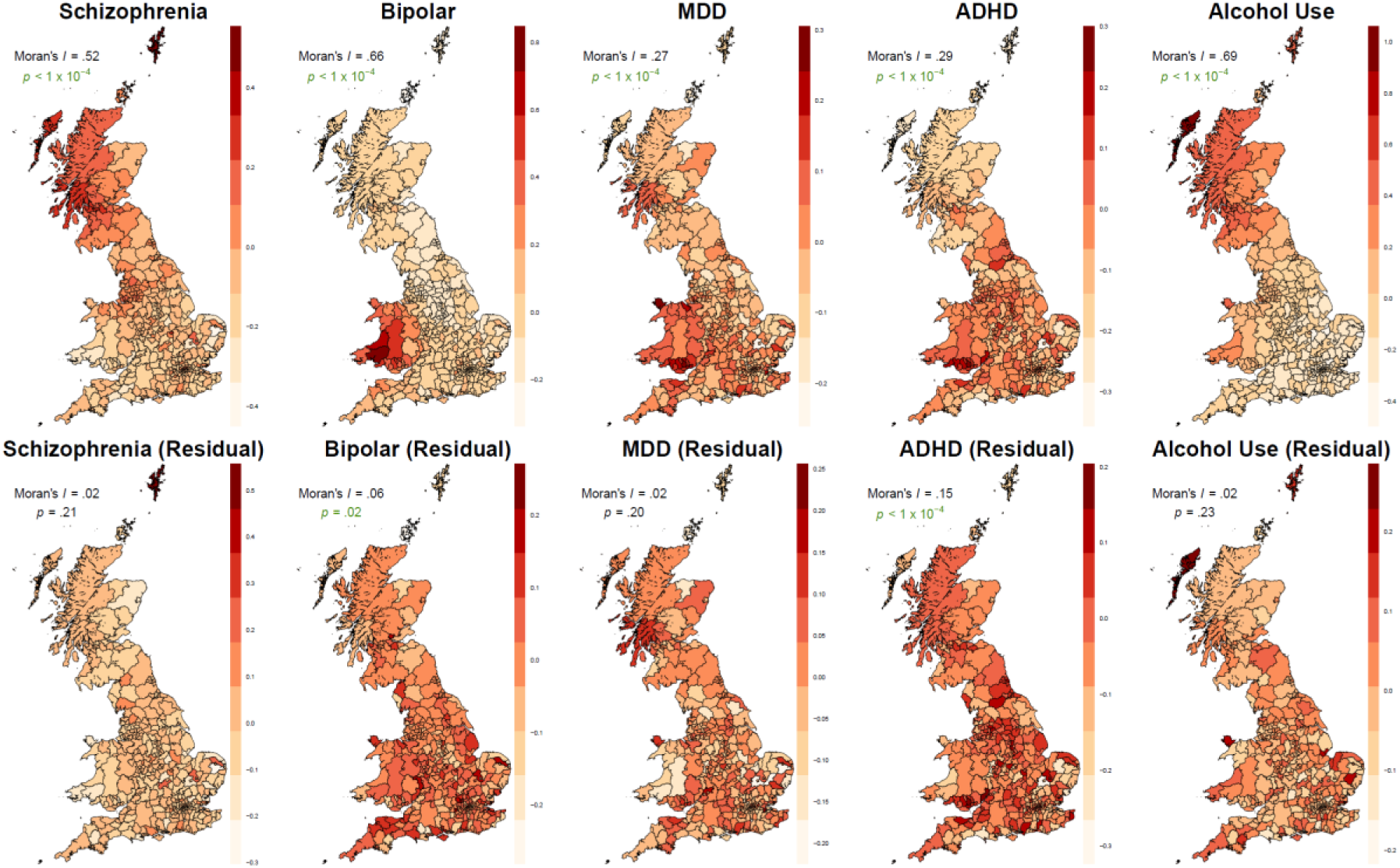
Geographic distribution (birthplace) and Moran’s *I* values for polygenic scores of four major psychiatric disorders (based on GWASs from the Psychiatric Genomics Consortium (PGC): schizophrenia^28^, bipolar^29^, MDD^30^, and ADHD^31^) and alcohol use^32^ before (top row) and after (bottom row) regressing out 100 ancestry-informative PCs. Green *p*-values are significant after FDR correction.

**Figure 3:**
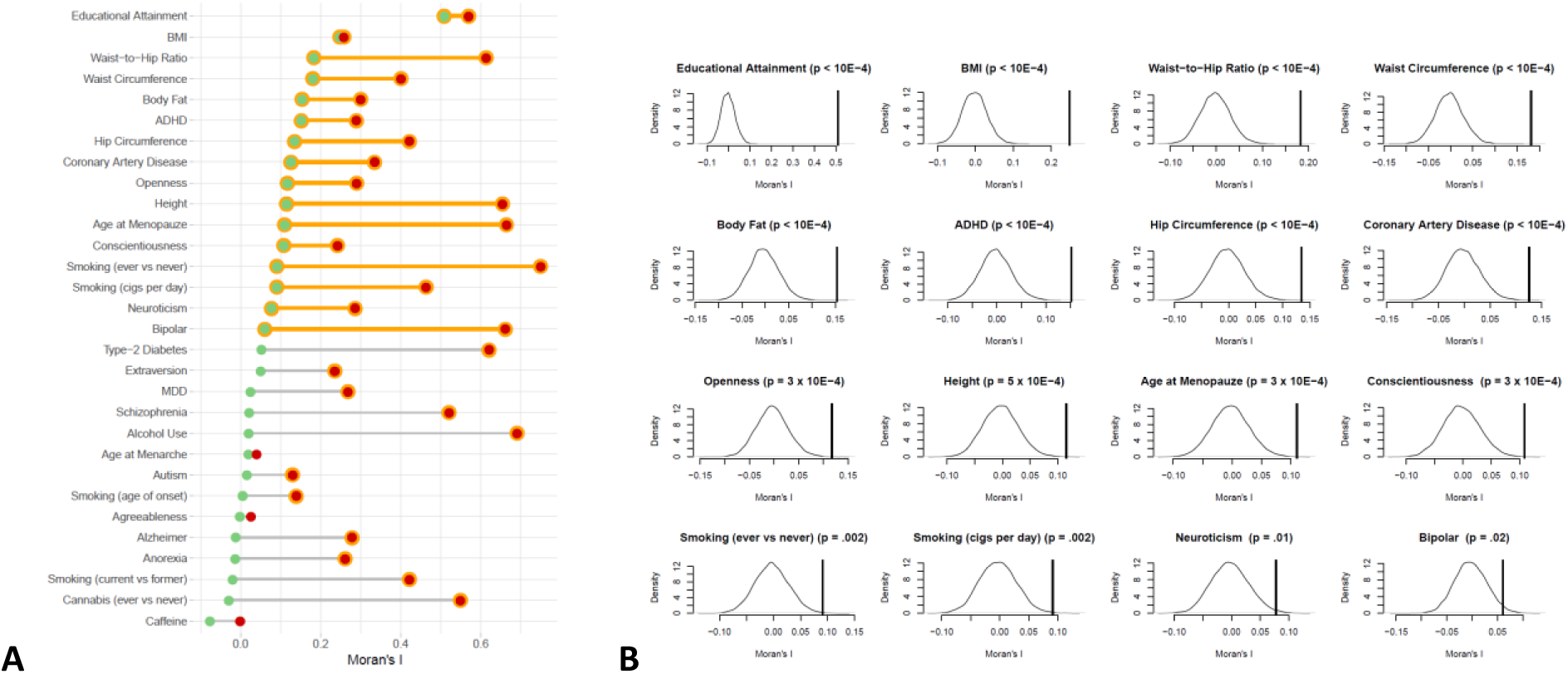
Moran’s *I* of 30 SBLUP polygenic scores computed using the average polygenic score per region in 378 local authority regions. **A** shows the Moran’s *I* of the polygenic scores unadjusted for PCs (red) and adjusted for 100 PCs (green), where orange means a significant FDR corrected *p*-value < .05 (corrected for 30 tests). **B** shows the distribution of significant Moran’s *I* statistics from 10,000 permutations that were conducted to obtain an empirical *p*-value for Moran’s *I*. The vertical line to the right of the permutation distribution shows the observed Moran’s *I* of the actual data.

**Figure 4:**
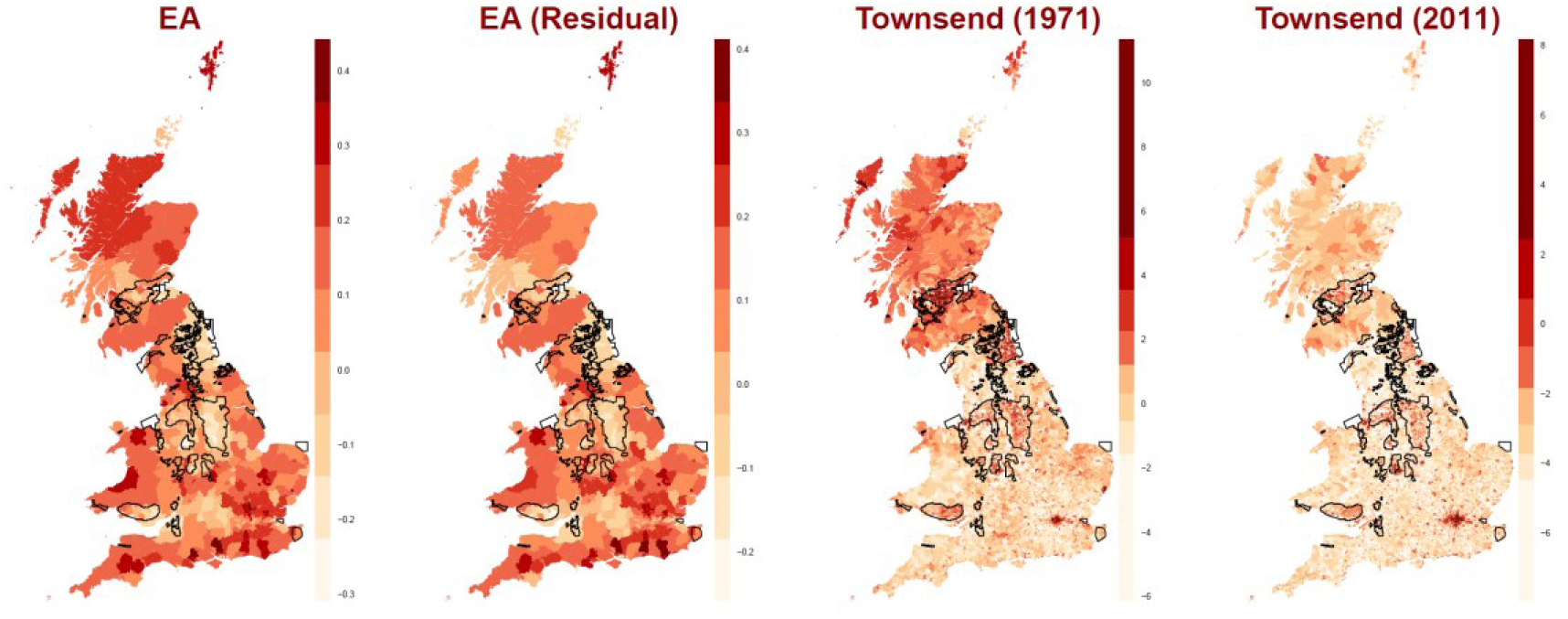
Geographic distribution (birthplace) of the educational attainment (EA) polygenic scores before and after regressing out 100 PCs, and the geographic distribution of Townsend indices from 1971 and 2011. The black lines indicate coal mining areas.

It has been argued that geographic clustering of complex trait genetic variation in UK Biobank is due to (subtle) ancestry differences or ascertainment bias.^27^ We discuss in more detail in the Supplementary Material why these are unlikely to be the sole explanations of our observations (paragraph: *Population Stratification and Ascertainment Bias*). In the next paragraph, we explore the more likely explanation, namely recent internal SES-related migrations.

## Consequences of SES-Related Migration

The geographic clustering of genome-wide trait-associated alleles after correcting for 100 PCs possibly reflects migration events that occurred more recently than the pre-modern demographic events that drove the regional ancestry differences captured by the PCs. Given the exceptionally strong geographic clustering of the EA score, we investigate here whether it reflects relatively recent internal migrations due to SES-related factors, which are known to especially motivate longer distance moves.^33^ Two types of migration flows may have affected the geographic clustering of SES-related alleles: 1) laborers and farmers leaving the country-side during the Industrial Revolution to work in the geographically clustered industrial jobs,^10^ and 2) more recent migration of higher-educated people, or people seeking a higher education, out of the more economically deprived industrial regions.

Much of the energy necessary for the mass-production that characterized the birth of the Industrial Revolution came from coal mines. The presence of coal and iron ore attracted large numbers of manual laborers. The Industrial Revolution and the later deindustrialization had a great impact on the economy of the coal mining areas.^34^ The decline of the British coal industry began in the 1920s, and nearly the whole industry has closed since the early 1980s, resulting in major job losses that still remain visible in unemployment rates decades later.^35^ Economic deprivation is widespread in coal mining areas: 43% of neighborhoods from coal mining areas fall into the 30% most economically deprived.^34^ In our analysis, coal mining areas show more economic deprivation than the rest of Great Britain from 1971 to 2011 as measured with the Townsend index^36^ (higher Townsend = more economic deprivation; all FDR corrected *p*-values < 10^−32^; see Figure 4 & Supplementary Figure 6). All regions have become less economically deprived over time, but the difference between coal mining areas and the rest remains highly significant.

After correcting for ancestry differences (100 PCs), the Townsend index is significantly associated with the 16 geographically clustered polygenic scores, with the strongest associations for EA (higher EA polygenic score = lower Townsend index; see Supplementary Figures 7 & 8). These 16 polygenic scores also all show significant differences between coal mining areas and the rest of the regions, both based on birth place and current address (Supplementary Figure 9), with EA showing the strongest differences (FDR corrected *p*-value < 10^−200^). We further compared ancestry-corrected polygenic scores between four groups of unrelated individuals: 1) people born in coal mining areas who moved out of coal mining areas (N=35,024), 2) people born outside of coal mining areas and still live outside of coal mining areas (N=129,298), 3) people born outside of coal mining areas who moved into coal mining areas (N=47,505), and 4) people born in coal mining areas who still live in coal mining areas (N=111,838). ANOVAs for all 16 geographically clustered polygenic scores show significant differences between the four groups (Figure 5), with EA showing the largest and most significant differences (*F*_3,323661_ = 687.3, FDR corrected *p*-value < 10^−200^). The largest differences were between people born in coal mining areas who moved away versus those who remained in the coal mining areas. The people that moved away have significantly higher EA polygenic scores than all other groups combined (t_43923_ = 19.8, *p* = 9 × 10^−87^), while those that remained have significantly lower EA polygenic scores than all other groups combined (t_230220_ = 44.6, *p* < 10^−200^). The degree of geographic clustering of polygenic scores, as measured by Moran’s *I*, is significantly correlated with the strength of their associations with Townsend, coal mining areas, and migration groups; the strongest correlations were between Moran’s *I* and the *F* statistic of the migration group differences: *r* = .95, *p* = 2 × 10^−8^ including EA and *r* = .73, *p* = .002 excluding EA (Supplementary Figure 10).

**Figure 5:**
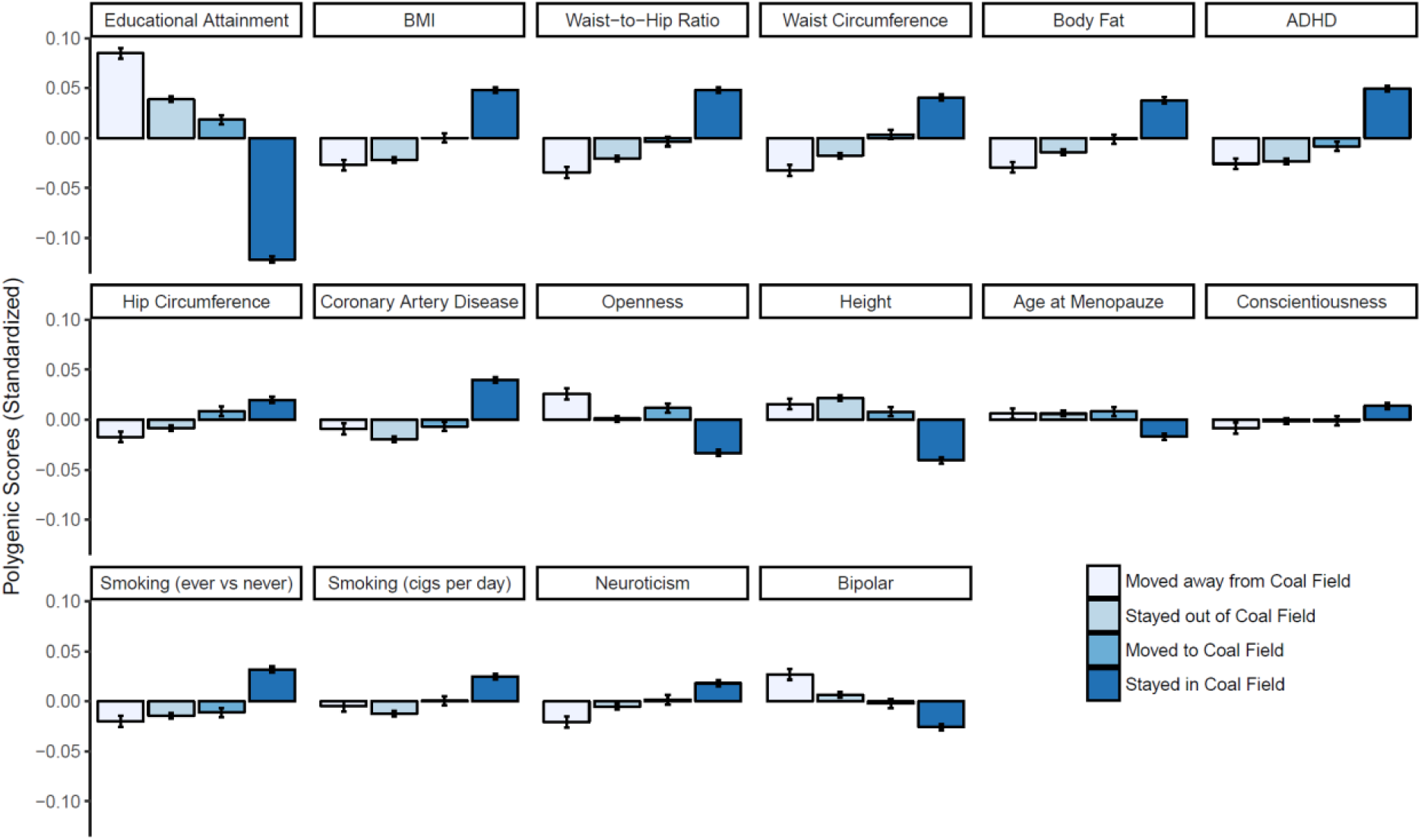
The average and standard errors of the 16 geographically clustered polygenic scores (ordered by Moran’s *I*) for four migration groups: born in coal field area and moved out, born in coal mining area and stayed, born outside of coal mining area and moved to coal mining area, born outside of coal mining area and stayed out. All polygenic scores shown are standardized residuals after regressing out 100 ancestry-informative PCs. ANOVAs were conducted for every polygenic score to test the presence of group differences, which were all significant with the least significant FDR corrected *p*-value of 1 × 10^−4^ for conscientiousness.

To get a better sense of the scale of regional differences in polygenic scores, and of how these change due to migration, we computed how much of the individual differences are explained by regional differences for both birthplace and current address (Supplementary Figure 15). The regional differences are greatest for the EA polygenic score, with about ~0.6-2.6% of individual differences being explained by regional differences, depending on how fine the regional scale is (the finer the scale, the more individual differences explained) and by whether the calculations are based on the birthplace or the current address. Regional differences are ~38-54% greater for the current address than for birthplace. The increase in variation explained by regional differences for the EA score (i.e., difference between birthplace and current address in % variance explained by region) is greater than the total variance explained by region for any other polygenic score. As would be expected from recent migration events, ancestry shows the opposite effect: comparing birthplace to current address, the variance explained by region has on average *decreased* by 37-73% for the first 30 PCs (Supplementary Figure 16).

## Genome-Wide Association Studies on Regional Outcomes

The geographic clustering of socio-economic resources and associated genetic variants may coincide with a range of regional collective views and attitudes. We examined this by leveraging the geographic clustering of genome-wide complex trait variation with GWASs on regional socio-economic and cultural outcomes, whereby all participants from the same region were assigned the same regional value as a phenotype (from here-on referred to as regional GWASs). The regional GWASs were run on the >400,000 UK Biobank participants, corrected for relatedness, age, sex, and ancestry (100 PCs). We first verified whether the approach works by running a regional GWAS on a regional measure of educational attainment (EA), obtained from census data, which showed genetic signals almost identical to an individual-level EA GWAS that excluded UK Biobank^37^ (see Supplementary Materials). We then ran regional GWASs on the presence of coal in an area and regional measures of major ideological factors known to cluster geographically, namely religiousness^11^ and political preference^12^. The regional socio-economic and cultural outcomes were defined as follows: whether the individual was born/lives in a coal mining area, the proportion of religious vs non-religious inhabitants in their region (based on current address), the proportion of “Leave” votes and non-voters in the 2016 Brexit referendum (current address), the proportion of non-voters and votes in the individuals’ constituency for three major UK parties in the UK 1970 general elections (based on birthplace) and the five major UK parties in the UK 2015 general elections (current address). We used the genome-wide summary statistics of the regional GWASs to estimate genetic correlations with a wide range of complex traits using LD score regression, a method that computes genetic correlations based on GWAS summary statistics without bias from sample overlap or ancestry differences.^22^ We summarize the main results on their (genetic) relationship with other complex traits below, and additional results in the Supplementary Materials.

The regional outcomes showed striking and often highly significant genetic correlations with a wide range of other traits (Figure 6). EA, IQ, and age at first birth showed significant genetic correlations with every regional outcome. Overall, the strongest genetic correlations were observed for cognition & SES-related traits (IQ, EA, income, and Townsend). These suggest that the election outcomes can be divided roughly into higher SES and lower SES regions, with Green Party, Liberal Democrats, and Conservative regions containing more alleles associated with higher SES trait values, and the Labour Party, UKIP, “Leave” votes for Brexit, and non-voters reflecting regions with more alleles associated with lower SES trait values. The election outcomes that are genetically associated with higher cognition and SES outcomes generally also show negative genetic correlations with disease risk outcomes (ADHD, MDD, smoking, alcohol dependence, heart disease, type-2 diabetes, BMI, longevity, and self-rated health), except for alcohol consumption, cannabis use, autism, and psychiatric disorders that are characterized by delusions (schizophrenia, bipolar, anorexia), for which the genetic association is the other way around (higher SES = higher risk). The genetic correlations were largely similar between election outcomes and the coal mining regions, likely due to the same systematic regional SES differences. A different pattern was observed for the proportion of religious inhabitants, which showed weaker genetic correlations with the SES related traits (cognition and health) and stronger associations with the two personality dimensions that showed significant geographic clustering in our previous analyses, openness and conscientiousness (more religious people = lower openness and higher conscientiousness). Risk taking, schizophrenia, and autism also show the highest genetic correlations with being religious (more religious people = lower genetic risk).

**Figure 6:**
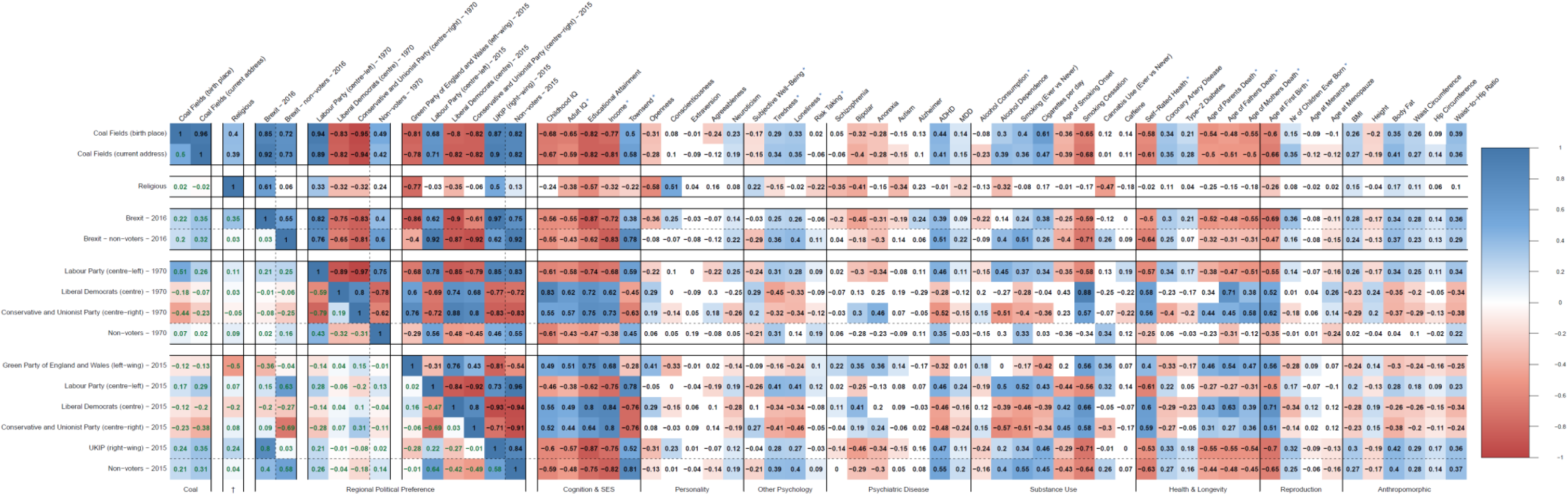
Genetic correlations based on LD score regression. Colored is significant after FDR correction. The green numbers in the left part of the Figure below the diagonal of 1’s are the phenotypic correlations between the regional outcomes of coal mining, religiousness, and regional political preference. The blue stars next to the trait names indicate that UK Biobank was part of the GWAS of the trait. See Supplementary Figure 23 for the standard errors and Supplementary Table 4 for the list of GWASs that the summary statistics of the complex traits were derived from.

The signals were largely consistent within parties over time (1970 & 2015) with respect to SES, but UKIP and Green Party did not yet exist in 1970. The genetic correlations between regional religiousness and the 1970 & 2015 election outcomes suggest that UKIP regions include former Labour Party regions with a more religious genetic profile (lower openness, higher conscientiousness), while the Green Party regions include former Conservative regions with a more non-religious genetic profile (higher openness, lower conscientiousness).

The genetic correlations between the regional outcomes were much stronger than the phenotypic correlations, and in some instances in opposite directions: Labour Party 2015 & Green Party (negative genetic, positive phenotypic correlation), Conservative Party 2015 & Green Party (positive genetic, negative phenotypic correlation), Conservative Party 2015 & Brexit (negative genetic, positive phenotypic correlation). A possible explanation is that the part of the regional variation that is explained by genetic differences is mostly related to regional socio-economic status (lower SES associated alleles in Labour and Brexit areas, higher SES in Conservative and Green areas), while environmental factors, which are responsible for most of the regional variation, are more characterized by ideology (Labour and Green areas being more left-wing, Conservative and Brexit more right-wing).

## Discussion

Understanding the consequences of DNA variation in human populations is of major importance for medical, biological, forensic, behavioral, and anthropological research. Since we have been able to measure DNA at a sequence level, studies have shown that the geographic distributions of alleles are not random and have mapped striking geographic patterns of ancestry.^2,3,5,38^ Here, we investigated geographic patterns of genome-wide complex trait variation and show that there are additional levels of genetic geographic clustering beyond the geographic patterns that reflect older ancestry differences. We show that the geographic clustering of genome-wide trait-associated alleles is related to recent geographic movement of people and that the resulting regional genetic patterns are associated with regional socio-economic and cultural outcomes.

Without controlling for ancestry, almost all traits we examined showed significant geographic clustering, often resembling the geographic patterns of ancestry differences within Great Britain. This indicates that either 1) the allele frequencies were differentiated between the different ancestries due to genetic drift or natural selection, and/or 2) the GWASs that produced the SNP effect estimates did not sufficiently control for ancestry differences, resulting in SNP effect estimates that are biased towards certain ancestral backgrounds. When we control for ancestry, 16 polygenic scores remain significantly clustered by geography. The strongest clustering was observed for EA. Among the rest of the geographically clustered traits are body dimensions, personality dimensions, and physical and mental health traits, which may reflect independent influences of them on non-random migration, and/or clustering that is (partly) driven by a genetic overlap with EA. The geographic clustering of complex trait variation seems to have increased due to relatively recent migration which is disrupting the older geographic patterns of ancestry (Supplementary Figures 15 & 16).

The degree of geographic clustering of the polygenic scores is largely in line with the strength of their relationship with regional economic deprivation and migration out of economically deprived regions (Supplementary Figure 10). People are more likely to migrate to improve their skills or employment prospects than for other area characteristics.^9^ Many industrialized countries showed these types of migration flows during the late 19^th^ and early 20^th^ century, where poorer laborers and small farmers left the country-side to work in industrial jobs that were often highly clustered in geographic space (e.g., coal mining areas).^12^ After the deindustrialization, the dense, durable, and affordable working-class houses and the public transportation networks from the industrial revolution remained in these neighborhoods and continued to attract poorer immigrants.^12^ Our results show that people with a genetic predisposition for higher cognitive abilities are leaving these regions, likely attracted by better educational or occupational opportunities in other regions. In fact, the people who were born in coal mining areas and migrated to better neighborhoods have higher average EA polygenic scores than people born outside of these regions. The regional clustering of cognitive abilities that follows may further affect the economic development of neighborhoods. These demographic processes may influence GWAS signals as well, where alleles that increase the chances of living in the unhealthy circumstances of lower SES neighborhoods may become part of the signal of a GWAS for a trait like BMI or body fat. There are for example significantly more McDonald’s restaurants in lower SES neighborhoods in Great Britain.^39^ This may be part of the explanation for why four out of the top five geographically clustered polygenic scores are related to body weight.

Selective migration has led to geographic clustering of social and economic needs, which can coincide with collective attitudes towards how communities should be organized and governed. We successfully captured heritability signals for regional religiousness and regional political attitudes, both of which have been shown to be partly heritable on an individual level^40–45^ and to cluster geographically^11,12^. From a regional genetic perspective, the election outcomes can be roughly divided into lower SES and higher SES electorates. Our findings suggest that the previously reported heritability estimates of these traits on an individual level may contain genetic effects on traits, such as EA, that influence which socio-economic strata and geographic regions people end up living in. Regional religiousness shows higher genetic correlations with personality (openness and conscientiousness) and less with the SES and health traits than the political parties do, which implies additional dimensions of geographic clustering beyond high versus low SES.

Our findings may largely reflect genetic consequences of social stratification, a key characteristic of human civilizations whereby society groups their people into strata based on SES. SES is generally based on occupation, income, and educational attainment, which are influenced by many environmental and genetic factors, and are associated with a wide range of physical and mental health outcomes. Socioeconomic status is not distributed randomly across geographic space, which leads to geographic clustering of alleles that are associated with SES-related traits such as educational attainment. Educational attainment is known for its high levels of assortative mating,^46,47^ which may be further induced by geographic clustering. This may affect social inequalities across generations through expanding biological inequalities in cognitive abilities and susceptibility to disease. It is possible that the combination of recent increases in social mobility and an improved educational system accelerates this separation of higher and lower genetic predisposition for traits related to cognition, SES, and health. Even though the genetic effects we find explain only part of the observed regional differences, researchers and social policy makers should keep these effects in mind, as they seem to be growing due to migration and can lead to detectable regional differences in health and social and economic success. For example, the significant genetic correlations between educational attainment and traits related to disease risk or body composition may decrease in the presence of stronger social safety nets that are geared towards making inhabitants of lower SES regions live more economically prosperous and healthier lives. Social policies that increase the quality of life in lower SES regions may also help to decrease migration out of the currently more economically deprived regions by people with genetic predispositions for higher SES outcomes, and thereby possibly result in a less geographically stratified society.

## Acknowledgements

This research was supported by the Australian National Health and Medical Research Council (1107258, 1078901, 1078037, 1056929, 1048853, and 1113400), and the Sylvia & Charles Viertel Charitable Foundation (Senior Medical Research Fellowship). B.P.Z. received funding from The Australian Research Council (FT160100298). M.G.N. is supported by ZonMw grants 849200011 and 531003014 from The Netherlands Organisation for Health Research and Development. K.J.H.V is supported by the Foundation Volksbond Rotterdam. This study makes use of data from the UK Biobank Resource (Application Number: 12514) and dbGaP (Accession Number: phs000674).

## Online Methods

### Participants

The participants of this study come from UK Biobank (UKB),^23^ which has received ethical approval from the National Health Service North West Centre for Research Ethics Committee (reference: 11/NW/0382). A total of 502,655 participants aged between 37 and 73 years old were recruited in the UK between 2006 and 2010. They underwent a wide range of cognitive, health, and lifestyle assessments, provided blood, urine, and saliva samples, and will have their health followed longitudinally.

### Genotypes and QC

A total of 488,377 UKB participants had their genome-wide single nucleotide polymorphisms (SNPs) genotyped on either the UK BiLEVE array (N = 49,950) or the UK Biobank Axiom Array (N = 438,423). The genotypes were imputed using the Haplotype Reference Consortium (HRC) panel as a reference set (pre-imputation QC and imputation are described in more detail in Bycroft et al, 2018).^48^ To create polygenic scores, we extracted a set of 1,312,100 autosomal HapMap 3 (HM3) SNPs with minor allele count (MAC) > 5, info score > 0.3, *p*_HWE_ < 10^−6^, and missingness < .05. For the genome-wide association study, we used 5.8 million SNPs that survived QC and have a MAF > .01.

### Ancestry & Principal Component Analysis

To capture British ancestry, we first excluded individuals with non-European ancestry. Ancestry was determined using Principal Component Analysis (PCA) in GCTA^49^. The UKB dataset was projected onto the first two principal components (PCs) from the 2,504 participants of the 1000 Genomes Project,^50^ using HM3 SNP with minor allele frequency (MAF) > 0.01 in both datasets. Next, participants from UKB were assigned to one of five super-populations from the 1000 Genomes project: European, African, East-Asian, South-Asian, or Admixed. Assignments for European, African, East-Asian, and South-Asian ancestries were based on > 0.9 posterior-probability of belonging to the 1000 Genomes reference cluster, with the remaining participants classified as Admixed. Posterior-probabilities were calculated under a bivariate Gaussian distribution where this approach generalizes the k-means method to take account of the shape of the reference cluster. We used a uniform prior and calculated the vectors of means and 2×2 variance-covariance matrices for each super-population. A total of 456,426 subjects were identified to have a European ancestry.

A PCA was then conducted on individuals of European ancestry in order to capture ancestry differences within the British population. In order to capture ancestry differences in homogenous populations, genotypes should be pruned for LD and long-range LD regions removed.^3^ The LD pruned (r^2^ < .1) UKB dataset without long-range LD regions consisted of 137,102 genotyped SNPs. The PCA to construct British ancestry-informative PCs was conducted on this SNP set for unrelated individuals using flashPCA v2.^51^ PC SNP loadings were used to project the complete set of European individuals onto the PCs.

### Polygenic Scores

Polygenic scores, the genome-wide sum of alleles weighted by their estimated effect sizes, were computed for 30 traits. The effect size estimates were obtained from genome-wide association studies (GWASs) that were chosen to not have included the UKB dataset to avoid over-estimation of the genetic predisposition of a trait.^25^ The polygenic scores were computed using the SBLUP approach,^46^ which maximizes the predictive power by creating scores with best linear unbiased predictor (BLUP) properties that account for linkage disequilibrium (LD) between SNPs. As a reference sample for the LD, we used the a random sample of 10,000 unrelated individuals from UK Biobank that were imputed using the Haplotype Reference Consortium (HRC) reference panel.^52^ The traits included psychiatric disorders, substance use, anthropomorphic traits, personality dimensions, educational attainment, reproduction, cardiovascular disease, and type-2 diabetes. Supplementary Table 1 lists the 30 traits and the GWASs from which we obtained the genome-wide effect sizes.

To further investigate the robustness of our results, we also created polygenic scores using only independent SNPs that were associated with the trait with a *p*-value < .05. The SNPs were clumped using PLINK^53^, using an r^2^ threshold of 0.1 and a window of 1 Mb as the physical distance threshold for clumping.

In order to examine the geographic clustering of polygenic scores beyond the clustering of ancestry, we created additional sets of polygenic scores that had the first 100 British ancestry-informative PCs regressed out.

### Spatial autocorrelations (Moran’s *I*)

The geographic clustering of ancestry and of genome-wide complex trait variation was investigated by testing whether the spatial autocorrelation (Moran’s *I*) is significantly greater than zero for ancestry-informative principal components (PCs), polygenic scores, and the residuals of polygenic scores after regressing out 100 ancestry-informative PCs. The spatial autocorrelation (Moran’s *I*) is the correlation in a measure among nearby locations in space, and its values range between −1 (dispersed) to 0 (spatially random) to 1 (spatially clustered).^26^ Moran’s *I*’s were computed using the average PCs or polygenic scores per region based on the birthplace of the subjects (378 regions, see Figure 1), whereby the regions were defined according to the local authorities division as provided by the UK Data Service InFuse database.^54^ The empirical *p*-values of Moran’s *I* statistics were derived with 10,000 permutations in which the average PCs or polygenic scores were permuted across regions (Figure 3B).

### Regional genome-wide association studies (GWASs)

Genome-wide association studies (GWASs) were run on publicly available regional outcomes, whereby all subjects from the same regions had the same regional phenotypic value assigned. Supplementary Figure 16 & 21 show the distributions of all phenotypes analyzed, except for the coal mining phenotypes, which were binary traits (47% of the participants were born in a coal mining area, and 50% of the participants currently live in a coal mining area). The regional phenotypes were obtained from the following public resources:

- The borders of a total of 208 coal mining regions were obtained from the *Coal Authority*: https://data.gov.uk/dataset/coal-mining-reporting-area
- The regional educational attainment for 342 local districts and for 7,195 Middle Super Output Areas (MSOA) was measured as the 2011 estimates of the highest qualification of residents of England >16 years old (5 levels: level 1 qualifications, level 2 qualifications, apprenticeship, level 3 qualifications, level 4 qualifications) was obtained from the *Nomis* database of the Office of National Statistics: https://www.nomisweb.co.uk/
- The proportion of religious vs non-religious inhabitants were obtained for 7,195 Middle Super Output Areas (MSOA) regions from the *Nomis* database of the Office of National Statistics: https://www.nomisweb.co.uk/
- The 2016 Brexit referendum results were obtained for 405 Local Authority Districts from *The Electoral Commision*: https://www.electoralcommission.org.uk/find-information-by-subject/elections-and-referendums/past-elections-and-referendums/eu-referendum/electorate-and-count-information
- The 1970 general election outcomes were obtained for 630 constituencies from *Political Science Resources*: http://www.politicsresources.net/area/uk/ge70/ge70index.htm
- The 2015 general election outcomes were obtained for 633 constituencies from *data.parliament.uk*: http://www.data.parliament.uk/dataset/general-election-2015 All political parties were included that had a median proportion of votes > 0.

We ran linear mixed model (LMM) GWASs with BOLT-LMM^55^ on participants with European ancestry, which controls for cryptic relatedness and population stratification by including a genetic relatedness matrix (GRM) in the model.^56^ Sex and age were included as covariates, as were the first 100 PCs as an additional control for population stratification. The results revealed a considerable inflation of test statistics that was not due to polygenic effects (this was captured by the LD score intercepts^57^ shown in Supplementary Table 2). This is likely due to the fact that participants that share regional environmental influences, because they come from the same region, are all assigned the same phenotypic value. We controlled for this inflation with an LD score intercept-based genomic control,^57^ i.e., we adjusted the standard errors (SE) of the estimated effect sizes as follows: 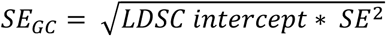 (see Supplementary Table 2).

### LD Score Regression

We partitioned the polygenic contributions to the heritability across genomic regions associated with histone modifications specific to ten cell-type/tissue groups using stratified LD score regression^58^ (Supplementary Figure 23). Genetic correlations were also computed using LD-score regression (Figure 6).^58^ The genetic correlation between traits is based on the estimated slope from the regression of the product of z-scores from two GWASs on the LD score and represents the genetic covariation between two traits based on all polygenic effects captured by the included SNPs. The genome-wide LD information used by these methods were based on European populations from the HapMap 3 reference panel.^57,58^ All LD score regression analyses included the 1,290,028 million genome-wide HapMap SNPs used in the original LD score regression studies.^57,58^

Computing genetic correlations with LD score regression is robust to sample overlap, so we included summary statistics from GWAS studies that also included UK Biobank (denoted with a blue star in Figure 6). Where possible however, we decided to display results obtained from summary statistics without UK Biobank, even if the GWASs from the original studies included UK Biobank participants. This was the case for MDD^30^ and educational attainment^37^, for which we used the same summary statistics that we used for the polygenic scores, namely from the GWASs that were re-run excluding UK Biobank. The genetic correlations for MDD and educational attainment obtained with the summary statistics that did include UK Biobank however were almost identical.

## Genome-Wide Association Studies on Regional Outcomes (Additional Results)

In order to empirically validate the approach, we first ran regional GWASs on regional measures of average EA outcomes as obtained from census data, namely the weighted average of 2011 estimates of the highest qualification of residents >16 years old, obtained from the Office for National Statistics (Supplementary Figure 16). This resulted in genetic signals that were very close to those of an individual-level EA GWAS that excluded UK Biobank.^37^ Most significant SNPs in the regional GWAS were at least nominally significant in the individual level GWAS and their effect sizes correlate .93 (Supplementary Figure 18). The genetic correlation between the regional EA GWAS and the individual level EA GWAS was .90, and the genetic correlations with 64 other complex traits were almost identical between the regional and individual level EA GWASs (*r* = .99, Supplementary Figures 19 & 20).

For the regional GWASs conducted on the presence of coal in the area, religiousness, and political preference, there were a total of 12 independent SNPs with *p* < 5 × 10^−8^ and 5 independent SNPs with *p* < 1 × 10^−8^ (Supplementary Figure 22 & Supplementary Table 3). The variance that could be accounted for by all SNPs (i.e., SNP heritability) ranged from 0.3% to 2.4% (see Supplementary Table 2), with the highest (≥2%) observed for Brexit, Green Party, UKIP, and non-voters in 2015. The heritability signals were significantly enriched for genetic variants that are active in hormonal pathways for the Green Party, and in the central nervous system for the Green Party, UKIP, 2015 non-voters, and Brexit (Supplementary Figure 23).

Regions with more non-voters genetically show a lower SES profile (i.e., strong negative genetic correlations with cognition and SES-related traits) and the largest positive genetic correlations with regions with more Labour party votes, up to .96 between the 2015 non-voters and the 2015 Labour voters. The 2015 non-voters regional GWAS shows the highest SNP heritability of the non-voters GWASs (2.2%). The genetic correlations also imply that regions with more non-voters and Labour voters show more risk-increasing alleles for mood-related traits (i.e., more MDD, higher neuroticism, more loneliness, and lower wellbeing), and no significant genetic correlation with conscientiousness, as opposed the other lower SES regions with more votes for UKIP and “Leave” votes for Brexit, which show a significant positive genetic correlation with conscientiousness.

In order to further examine what differentiates the parties within the higher SES and lower SES clusters from each other, we repeated the regional GWASs for the proportion of votes among only the Green Party, Liberal Democrats, and Conservatives votes, and the proportion of votes among only Labour Party and UKIP votes. The correlations with religiousness were consistently higher and more often significant for the differences within the higher and lower SES voters than the differences between them (Supplementary Figure 24). The genetic signals that differentiate Green Party regions from the other higher SES votes show the highest genetic correlation with regional religiousness (more Green Party votes = less religious: *r*_g_ = -.82, SE = .06). What differentiates Liberal Democrats from the other higher SES parties still seems to be largely SES-related, given the high positive genetic correlation with EA and income (both .77). The lower SNP heritability estimates of the within SES differences (0.3% - 1.2%, compared to 1.1% - 2.4%, see Supplementary Table 2) suggest that the differences within the higher and lower SES voters are less influenced by regional genetics than the differences between them.

## Population Stratification and Ascertainment Bias

### Population Stratification

The largest patterns of genome-wide variation between and within human populations are due to differences in ancestry rather than trait variation. In genetic association studies, false positives due to population stratification occur when these systematic ancestry differences get mistaken for associations due to genetic variants that influence the trait that is being studied.^59^ False positives due to population stratification can occur when trait differences are in line with ancestry differences, which could also occur due to non-genetic factors, such as regional differences in environmental exposures. Geographic location is known to strongly correlate with ancestry differences: the closer people live to each other, the more likely it is that they share more ancestors. The main focus of this study is the relationship between geographic location and genome-wide complex trait variation, which is why we had to be particularly rigorous in accounting for population stratification. We summarize below why it is unlikely that our observations are merely a result of ancestry differences or biased polygenic scores.

- The most widely used approach to account for ancestry differences is to quantify ancestry differences with a principal component analysis (PCA) on genome-wide SNP data and then account for the resulting principal components (PCs).^24,59^ Instead of using the standard 40 PCs provided by UK Biobank, which capture both non-European and European ancestry differences,^48^ we re-computed PCs to more effectively capture population stratification within the more homogeneous group of British participants with European ancestry (see Online Methods). While genome-wide association studies (GWASs) usually control for 10 to 40 PCs, we controlled for the first 100 PCs in all our analyses.
- We validated the effectiveness of the 100 PCs in accounting for geographic clustering due to population stratification using polygenic scores that reflect European ancestry differences as captured in an independent European-American dataset: the GERA cohort.^60^ First, we conducted GWASs in GERA (N = 51,258) on the first 20 GERA PCs in order to get SNP effects that reflect genome-wide patterns of their European-American ancestry differences. We then used these SNP effects to build polygenic scores in UK Biobank. These all show significant geographic clustering as quantified with Moran’s *I*. After controlling for 100 PCs from UKB all Moran’s *I*’s dropped to being not significantly greater than 0 (see Online Methods and Supplementary Figures 3 and 4).
- The polygenic scores we analyzed were constructed from 1,312,100 autosomal SNPs, regardless of how significantly associated they are with the trait. The ensemble of non-significant SNPs contain a substantial amount of signals due to true causal relationships, and thus meaningful effect sizes, which increase the predictive power of polygenic scores.^61^ Increasing the number of non-associated SNPs however may also increase the chances of including more stratified SNPs in the polygenic score. We therefore created a set of additional polygenic scores using only independent SNPs (i.e., clumped) that were at least nominally significantly associated with the trait at *p* < .05. This results in scores that are based on fewer SNPs that are more reliably associated with the trait, but also results in less predictive scores. With this approach, 8 out of 16 previously significant traits are significantly geographically clustered after FDR correction (see Supplementary Figure 2). These geographically clustered clumped scores also showed similar and significant associations with Townsend (Supplementary Figures 11 & 12), coal mining regions (Supplementary Figure 13) and migration out of coal fields (Supplementary Figures 14), with educational attainment showing the strongest effects.
- We show that the geographically clustered polygenic scores are significantly associated with regional outcomes of economic deprivation and with migration out of the more economically deprived regions in the UK (coal mining regions). The strength of the geographic clustering is in line with the strength of the association with regional outcomes and, more importantly, migration (Supplementary Figure 10). In other words, 1) the traits that show significant geographic clustering are the traits that cluster in specific regions that are characterized by lower SES measures, and 2) we show that the processes that would result in these regional differences are measurable in the current dataset, namely migration out of the lower SES regions by individuals with a higher predisposition for SES-related traits such as higher educational attainment and lower body weight. Although these observations do not directly prove that ancestry differences cannot account for this geographic clustering, it does show that *if* subtle population stratification would be the cause of these regional differences (which is unlikely given our stringent control for ancestry differences), it would have to involve ancestry differences that are in line with genome-wide complex trait variation.
- SNPs that are in LD with many SNPs are more likely to tag a causal SNP (i.e., be correlated with a causal SNP), and are thus more likely to have a higher test-statistic in a GWAS. The amount of SNPs that is tagged by a SNP is quantified by its LD Score. LD Score regression is an approach that leverages the relationship between the LD score of a SNP and the GWAS test statistic to distinguish inflation of genome-wide test statistics due to variants that influence the complex trait under study from inflation due to confounding bias such as population stratification.^57^ LD Score regression analyses show that the results from our regional GWASs all show an inflation of test statistics that is partly due to confounding (likely shared environmental influences) but also contains a considerable inflation due to variants that are associated with complex trait variation that is being captured with the regional measures. LD Score regression was then used to compare the parts of the genetic signals that were due to causal variants between our regional GWASs and GWASs from a wide range of other complex traits. Importantly, LD Score regression showed that the signals from our regional GWAS on EA contained almost the same signals as an individual level GWAS on EA that was conducted on non-UK Biobank datasets, which is in line with the geographic clustering of genome-wide alleles that have a causal influence on EA.

### Ascertainment Bias

The UK Biobank ascertainment strategy was designed to capture sufficient variation in socioeconomic, urban–rural, and ethnic background.^23^ The participation rate however was 5.45% and was biased towards older, more healthy, and female residents.^62^ The UK Biobank sample does reflect nationally representative data sources to a significant degree, making it likely that our observations would generalize to the population at large. We tested at the MSOA level how EA measurements in UK Biobank compare to nationally representative census data (the same EA census measurements that we used for the regional EA GWAS). The average EA per MSOA region as measured in UK Biobank is strongly predictive of MSOA-EA as measured from the nationally representative census data (*p* < 10^−16^, *R*^2^ = 40%; Supplementary Figure 5). The average polygenic scores per MSOA region, with 100 PCs regressed out, are also highly predictive of MSOA-EA according to nationally representative census data (*p* < 10^−16^, *R*^2^ = 19%; Supplementary Figure 5). Since UK Biobank has sampled healthier individuals as well as fewer individuals from more economically deprived areas as compared to the British population as a whole,^62^ the regional differences that we report may turn out to be stronger in the real population than in the UK Biobank sample.

## Supplementary Note: Acknowledgements

GERA: The Genetic Epidemiology Research on Adult Health and Aging study was supported by grant RC2 AG036607 from the National Institutes of Health, grants from the Robert Wood Johnson Foundation, the Ellison Medical Foundation, the Wayne and Gladys Valley Foundation and Kaiser Permanente. The authors thank the Kaiser Permanente Medical Care Plan, Northern California Region (KPNC) members who have generously agreed to participate in the Kaiser Permanente Research Program on Genes, Environment and Health (RPGEH).

UK Biobank: This study has been conducted using UK Biobank resource under Application Number 12514. UK Biobank was established by the Wellcome Trust medical charity, Medical Research Council, Department of Health, Scottish Government and the Northwest Regional Development Agency. It has also had funding from the Welsh Assembly Government, British Heart Foundation and Diabetes UK.

**Supplementary Table 1:** Overview of GWASs that provided the effect size estimates that were used to compute polygenic scores

| Trait Name | N<sub>GWAS</sub> | N<sub>cases</sub> | N<sub>controls</sub> | N<sub>effective</sub> | h<sup>2</sup><sub>SNP</sub> | LDSC Intercept |
| --- | --- | --- | --- | --- | --- | --- |
| Educational Attainment<sup>37</sup> | 217,569 | - | - | 217,569 | .10 | 1.02 |
| **Personality:** |  |  |  |  |  |  |
| Openness<sup>63,64</sup> | 76,551 | - | - | 76,551 | .10 | .95 |
| Conscientiousness<sup>63,64</sup> | 76,551 | - | - | 76,551 | .08 | .96 |
| Extraversion<sup>63,64</sup> | 76,600 | - | - | 76,600 | .14 | .92 |
| Agreeableness<sup>63,64</sup> | 76,548 | - | - | 76,548 | .07 | .96 |
| Neuroticism<sup>63,64</sup> | 76,581 | - | - | 76,581 | .09 | .94 |
| **Psychiatric Traits:** |  |  |  |  |  |  |
| Schizophrenia<sup>28</sup> | 150,064 | 36,989 | 113,075 | 111,486.6 | .18 | 1.05 |
| Bipolar<sup>29</sup> | 63,766 | 11,974 | 51,792 | 38,902.1 | .14 | 1.02 |
| Anorexia<sup>65</sup> | 14,477 | 3,495 | 10,982 | 10,605 | .22 | 1.01 |
| Autism<sup>66</sup> | 46,350 | 18,381 | 27,969 | 44,366.62 | .11 | 1.01 |
| Alzheimer<sup>67</sup> | 54,162 | 17,008 | 37,154 | 46,668.5 | .08 | 1.04 |
| ADHD<sup>31</sup> | 55,374 | 20,183 | 35,191 | 51,306.4 | .21 | 1.03 |
| MDD<sup>30</sup> | 431,394 | 116,404 | 314,990 | 161,089.68 | .08 | 1.01 |
| **Substance Use:** |  |  |  |  |  |  |
| Alcohol Use<sup>32</sup> | 70,493 | - | - | 70,493 | .05 | 1.02 |
| Smoking (ever vs never)<sup>68</sup> | 74,035 | 41,969 | 32,066 | 72,710.4 | .12 | 1.00 |
| Smoking (cigs per day)<sup>68</sup> | 38,181 | - | - | 38,181 | .06 | 1.01 |
| Smoking (current/former)<sup>68</sup> | 41,278 | 23,969 | 17,309 | 40,203.4 | .09 | 1.01 |
| Smoking (age at onset)<sup>68</sup> | 24,114 | - | - | 24,114 | .06 | 1.00 |
| Cannabis (ever vs never)<sup>69</sup> | 32,330 | 14,387 | 17,943 | 31,938.9 | .13 | 1.00 |
| Caffeine<sup>70</sup> | 91,462 | - | - | 91,462 | .04 | 1.01 |
| **CAD & Diabetes:** |  |  |  |  |  |  |
| Coronary Artery Disease<sup>71</sup> | 86,995 | 22,233 | 64,762 | 66,204 | .28 | .87 |
| Type-2 Diabetes<sup>72</sup> | 69,033 | 12,171 | 56,862 | 40,100.7 | .18 | 1.01 |
| **Reproduction:** |  |  |  |  |  |  |
| Age at Menarche<sup>73</sup> | 87,802 | - | - | 87,802 | .24 | .98 |
| Age at Menopause<sup>74</sup> | 69,360 | - | - | 69,360 | .13 | .99 |
| **Antropomorphic traits:** |  |  |  |  |  |  |
| Height<sup>75</sup> | 253,280 | - | - | 253,280 | .31 | 1.33 |
| BMI<sup>76</sup> | 322,154 | - | - | 322,154 | .13 | .67 |
| Body Fat<sup>77</sup> | 100,716 | - | - | 100,716 | .10 | .91 |
| Waist Circumference<sup>78</sup> | 232,101 | - | - | 232,101 | .13 | .76 |
| Hip Circumference<sup>78</sup> | 213,038 | - | - | 213,038 | .14 | .78 |
| Waist-to-hip-ratio<sup>78</sup> | 212,248 | - | - | 212,248 | .10 | .84 |

**Supplementary Table 2:** Sample sizes and LD score regression results for the regional GWASs before and after LDSC-intercept based genomic control.

|  |  | Before Genomic Control |  |  |  | After Genomic Control (LDSC intercept=1; ratio=0) |  |
| --- | --- | --- | --- | --- | --- | --- | --- |
| Regional Phenotype | N | SNP-h<sup>2</sup> (SE) % | lambda | LDSC intercept (SE) | ratio | SNP-h<sup>2</sup> (SE) % | lambda |
| 2011 - Educational attainment (local authorities) | 402,552 | 4.1 (.25) | 1.49 | 1.22 (.01) | .39 (.02) | 3.3 (.20) | 1.23 |
| 2011 - Educational attainment (MSOA) | 416,061 | 7.5 (.29) | 1.63 | 1.14 (.01) | .18 (.01) | 6.6 (.25) | 1.44 |
| Coal Fields (Birth Place) | 422,757 | 1.5 (.20) | 1.56 | 1.43 (.01) | .77 (.02) | 1.0 (.12) | 1.07 |
| Coal Fields (Current Address) | 449,972 | 2.1 (.18) | 1.43 | 1.25 (.01) | .55 (.02) | 1.7 (.13) | 1.14 |
| 2011 - Proportion of non-religious individuals | 416,061 | 1.2 (.17) | 1.31 | 1.19 (.01) | .67 (.03) | 1.0 (.13) | 1.08 |
| 2016 - Brexit | 446,910 | 3.0 (.22) | 1.56 | 1.31 (.01) | .52 (.02) | 2.3 (.16) | 1.19 |
| 2016 - Non-voters | 446,910 | 1.3 (.17) | 1.20 | 1.12 (.01) | .50 (.04) | 1.2 (.14) | 1.09 |
| 1970 - Labour Party (centre-left) | 422,189 | 2.3 (.22) | 1.49 | 1.30 (.01) | .60 (.02) | 1.8 (.15) | 1.14 |
| 1970 - Liberal Democrats (centre) | 207,045 | 0.9 (.30) | 1.15 | 1.12 (.01) | .75 (.05) | 0.8 (.25) | 1.04 |
| 1970 - Conservative and Unionist Party (centre-right) | 421,940 | 1.6 (.18) | 1.37 | 1.27 (.01) | .66 (.02) | 1.3 (.13) | 1.10 |
| 1970 - Non-voters | 422,415 | 0.6 (.17) | 1.31 | 1.28 (.01) | .83 (.02) | 0.5 (.12) | 1.04 |
| 2015 - Green Party of England and Wales (left-wing) | 449,553 | 2.2 (.16) | 1.25 | 1.08 (.01) | .27 (.03) | 2.0 (.14) | 1.18 |
| 2015 - Labour Party (centre-left) | 449,553 | 1.8 (.19) | 1.37 | 1.22 (.01) | .57 (.02) | 1.4 (.14) | 1.13 |
| 2015 - Liberal Democrats (centre) | 449,553 | 1.2 (.15) | 1.20 | 1.08 (.01) | .42 (.05) | 1.1 (.14) | 1.09 |
| 2015 - Conservative and Unionist Party (centre-right) | 449,553 | 1.5 (.17) | 1.31 | 1.17 (.01) | .56 (.03) | 1.2 (.14) | 1.11 |
| 2015 - UKIP (right-wing) | 449,553 | 3.1 (.21) | 1.56 | 1.29 (.01) | .50 (.02) | 2.4 (.15) | 1.20 |
| 2015 - Non-voters | 449,553 | 2.6 (.19) | 1.43 | 1.19 (.01) | .43 (.02) | 2.2 (.15) | 1.19 |
| 2015 - Green/(Green+Liberals+Conservative) | 449,553 | 1.2 (.14) | 1.15 | 1.06 (.01) | .31 (.04) | 1.2 (.13) | 1.11 |
| 2015 - Liberals/(Green+Liberals+Conservative) | 449,553 | 0.3 (.12) | 1.10 | 1.08 (.01) | .72 (.07) | 0.3 (.11) | 1.03 |
| 2015 - Conservative /(Green+Liberals+Conservative) | 449,553 | 0.9 (.13) | 1.15 | 1.08 (.01) | .51 (.05) | 0.8 (.11) | 1.07 |
| 2015 - Labour/(UKIP+Labour) | 449,553 | 0.7 (.14) | 1.20 | 1.15 (.01) | .70 (.04) | 0.6 (.11) | 1.05 |
| 2015 - UKIP/(UKIP+Labour) | 449,553 | 0.7 (.14) | 1.20 | 1.15 (.01) | .70 (.04) | 0.6 (.11) | 1.05 |

**Supplementary Table 3:** Independent SNPs with *p* < 5 × 10^−8^, with independence based on an *r*^2^ threshold of 0.1.

| Election Phenotype | SNP | Chr | BP (hg19) | p-value | MAF | Gene |
| --- | --- | --- | --- | --- | --- | --- |
| Coal Mining (Birth Place) | rs143252507 | 17 | 19970702 | $2.5 \times 10^{-8}$ | .01 | SPECC1 |
| 2016 - Brexit | **rs7613360** | **3** | **49916710** | **$9.3 \times 10^{-10}$** | **.40** | **ACTBP13** |
| | rs1050450 | 3 | 49394834 | $2.5 \times 10^{-8}$ | .31 | GPX1 |
| | rs6130360 | 20 | 42010996 | $3.1 \times 10^{-8}$ | .15 | |
| 1970 - Labour Party (centre-left) | rs7159181 | 14 | 39229004 | $1.4 \times 10^{-8}$ | .44 | LINC00639 |
| 2015 - Green Party of England and Wales (left-wing) | **rs2624838** | **3** | **50205642** | **$8.3 \times 10^{-10}$** | **.34** | **SEMA3F** |
|  | **rs13135092** | **4** | **103198082** | **$1.1 \times 10^{-9}$** | **.08** | **SLC39A8** |
| 2015 - Conservative and Unionist Party (centre-right) | rs61278749 | 10 | 106577845 | $3.0 \times 10^{-8}$ | .13 | SORCS3 |
| 2015 - UKIP (right-wing) | **rs9827708** | **3** | **49649989** | **$2.4 \times 10^{-11}$** | **.30** | **BSN** |
| | rs2624848 | 3 | 50165101 | $2.0 \times 10^{-8}$ | .44 | |
| 2015 – Non-voters | **rs7548936** | **1** | **91207757** | **$8.4 \times 10^{-11}$** | **.37** |  |
| | rs79003713 | 5 | 138135234 | $3.6 \times 10^{-8}$ | .06 | CAP102 |

*Bold: $p < 1 \times 10^{-8}$*

**Supplementary Table 4:** Overview of GWASs that provided the effect size estimates that were used to compute genetic correlations with regional GWASs using LD score regression.

| Trait Name | NGWAS | Ncases | Ncontrols | Neffective | h<sup>2</sup><sub>SNP</sub> | LDSC Intercept |
| --- | --- | --- | --- | --- | --- | --- |
| **Cognition and SES:** |  |  |  |  |  |  |
| Childhood IQ<sup>79</sup> | 12,441 | - | - | 12,441 | .28 | 1.00 |
| Adult IQ<sup>80</sup> | 78,308 | - | - | 78,308 | .19 | 1.02 |
| Educational Attainment<sup>37</sup> | 217,569 | - | - | 217,569 | .10 | 1.02 |
| Income<sup>81</sup> | 112,151 | - | - | 112,151 | .06 | 1.03 |
| Townsend<sup>81</sup> | 112,151 | - | - | 112,151 | .04 | 1.01 |
| **Personality:** |  |  |  |  |  |  |
| Openness<sup>63,64</sup> | 76,551 | - | - | 76,551 | .10 | .95 |
| Conscientiousness<sup>63,64</sup> | 76,551 | - | - | 76,551 | .08 | .96 |
| Extraversion<sup>63,64</sup> | 76,600 | - | - | 76,600 | .14 | .92 |
| Agreeableness<sup>63,64</sup> | 76,548 | - | - | 76,548 | .07 | .96 |
| Neuroticism<sup>63,64</sup> | 76,581 | - | - | 76,581 | .09 | .94 |
| **Other Psychology:** |  |  |  |  |  |  |
| Subjective Well-being<sup>82</sup> | 298,420 | - | - | 298,420 | .03 | 1.00 |
| Tiredness<sup>83</sup> | 108,976 | - | - | 108,976 | .07 | .99 |
| Loneliness<sup>84</sup> | 332,991 | - | - | 332,991 | .08 | 1.01 |
| Risk Taking<sup>84</sup> | 332,991 | - | - | 332,991 |  |  |
| **Psychiatric Traits:** |  |  |  |  |  |  |
| Schizophrenia<sup>28</sup> | 150,064 | 36,989 | 113,075 | 111,486.6 | .18 | 1.05 |
| Bipolar<sup>29</sup> | 63,766 | 11,974 | 51,792 | 38,902.1 | .14 | 1.02 |
| Anorexia<sup>65</sup> | 14,477 | 3,495 | 10,982 | 10,605 | .22 | 1.01 |
| Autism<sup>66</sup> | 46,350 | 18,381 | 27,969 | 44,366.62 | .11 | 1.01 |
| Alzheimer<sup>67</sup> | 54,162 | 17,008 | 37,154 | 46,668.5 | .08 | 1.04 |
| ADHD<sup>31</sup> | 55,374 | 20,183 | 35,191 | 51,306.4 | .21 | 1.03 |
| MDD<sup>30</sup> | 431,394 | 116,404 | 314,990 | 161,089.68 | .08 | 1.01 |
| **Substance Use:** |  |  |  |  |  |  |
| Alcohol Consumption<sup>85</sup> | 112,117 | - | - | 112,117 | .08 | 1.01 |
| Alcohol Dependence<sup>86</sup> | 46,568 | 11,569 | 34,999 | 34,779.54 | .18 | 1.02 |
| Smoking (ever vs never)<sup>68</sup> | 74,035 | 41,969 | 32,066 | 72,710.4 | .12 | 1.00 |
| Smoking (cigs per day)<sup>68</sup> | 38,181 | - | - | 38,181 | .06 | 1.01 |
| Smoking (current/former)<sup>68</sup> | 41,278 | 23,969 | 17,309 | 40,203.4 | .09 | 1.01 |
| Smoking (age at onset)<sup>68</sup> | 24,114 | - | - | 24,114 | .06 | 1.00 |
| Cannabis (ever vs never)<sup>69</sup> | 32,330 | 14,387 | 17,943 | 31,938.9 | .13 | 1.00 |
| Caffeine<sup>70</sup> | 91,462 | - | - | 91,462 | .04 | 1.01 |
| **Health & Longevity:** |  |  |  |  |  |  |
| Self-rated health<sup>87</sup> | 111,749 | - | - | 111,749 |  |  |
| Coronary Artery Disease<sup>71</sup> | 86,995 | 22,233 | 64,762 | 66,204 | .28 | .87 |
| Type-2 Diabetes<sup>72</sup> | 69,033 | 12,171 | 56,862 | 40,100.7 | .18 | 1.01 |
| Age of Parents Death<sup>88</sup> | 45,627 | - | - | 45,627 | .05 | 1.02 |
| Age of Fathers Death<sup>88</sup> | 63,775 | - | - | 63,775 | .05 | 1.02 |
| Age of Mothers Death<sup>88</sup> | 52,776 | - | - | 52,776 | .05 | 1.01 |
| **Reproduction:** |  |  |  |  |  |  |
| Age at First Birth<sup>89</sup> | 251,151 | - | - | 251,151 | .05 | .96 |
| Nr of Children Ever Born<sup>89</sup> | 343,072 | - | - | 343,072 | .02 | .97 |
| Age at Menarche<sup>73</sup> | 87,802 | - | - | 87,802 | .24 | .98 |
| Age at Menopause<sup>74</sup> | 69,360 | - | - | 69,360 | .13 | .99 |
| **Antropomorphic:** |  |  |  |  |  |  |
| BMI<sup>76</sup> | 322,154 | - | - | 322,154 | .13 | .67 |
| Height<sup>75</sup> | 253,280 | - | - | 253,280 | .31 | 1.33 |
| Body Fat<sup>77</sup> | 100,716 | - | - | 100,716 | .10 | .91 |
| Waist Circumference<sup>78</sup> | 232,101 | - | - | 232,101 | .13 | .76 |
| Hip Circumference<sup>78</sup> | 213,038 | - | - | 213,038 | .14 | .78 |
| Waist-to-hip-ratio<sup>78</sup> | 212,248 | - | - | 212,248 | .10 | .84 |

**Supplementary Figure 1:**
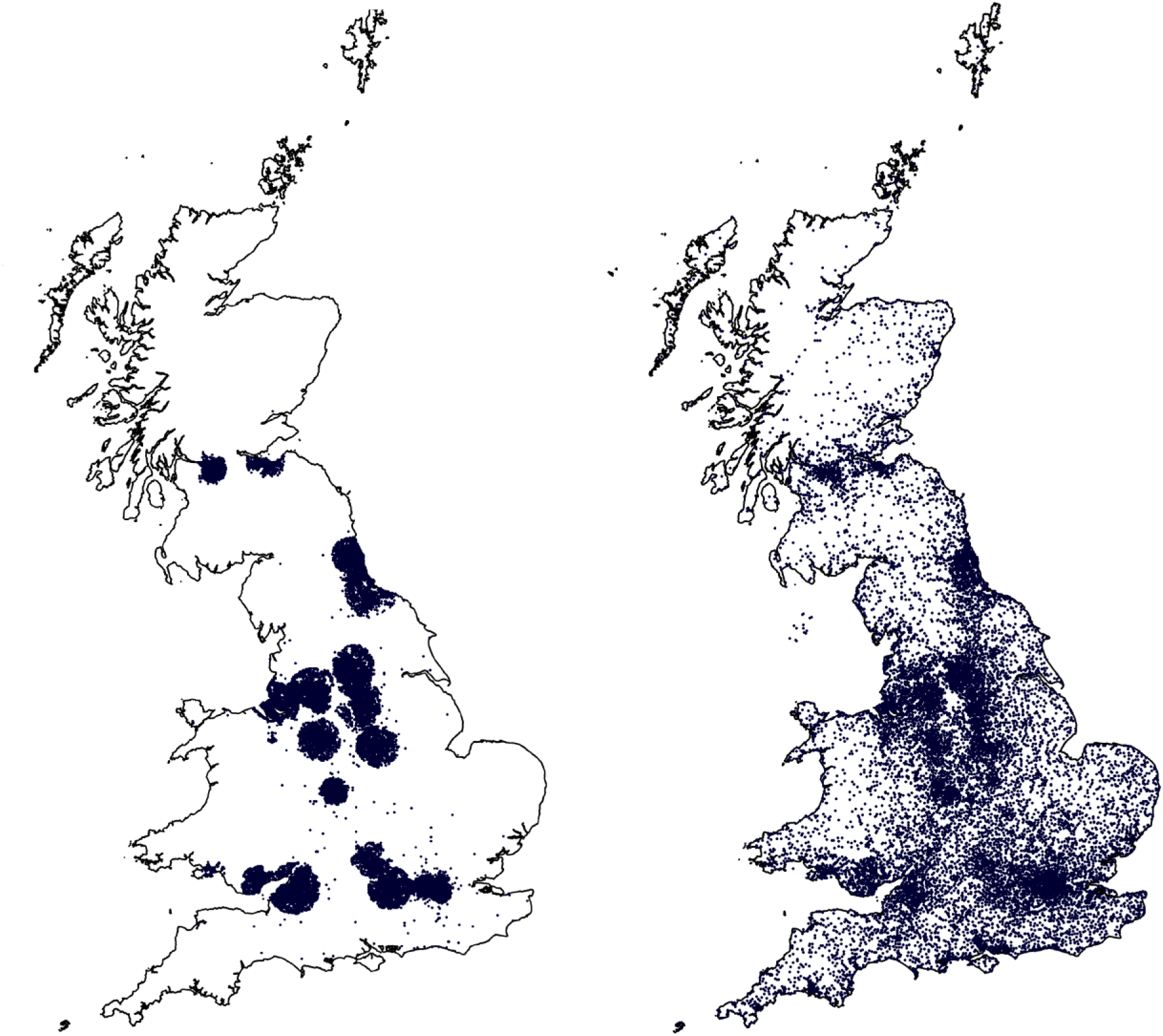
The geographic locations of the UK Biobank participants (each dot represents a participant). The left map shows the current living address (N=497,673). The right plot shows the birth places (N=444,992).

**Supplementary Figure 2:**
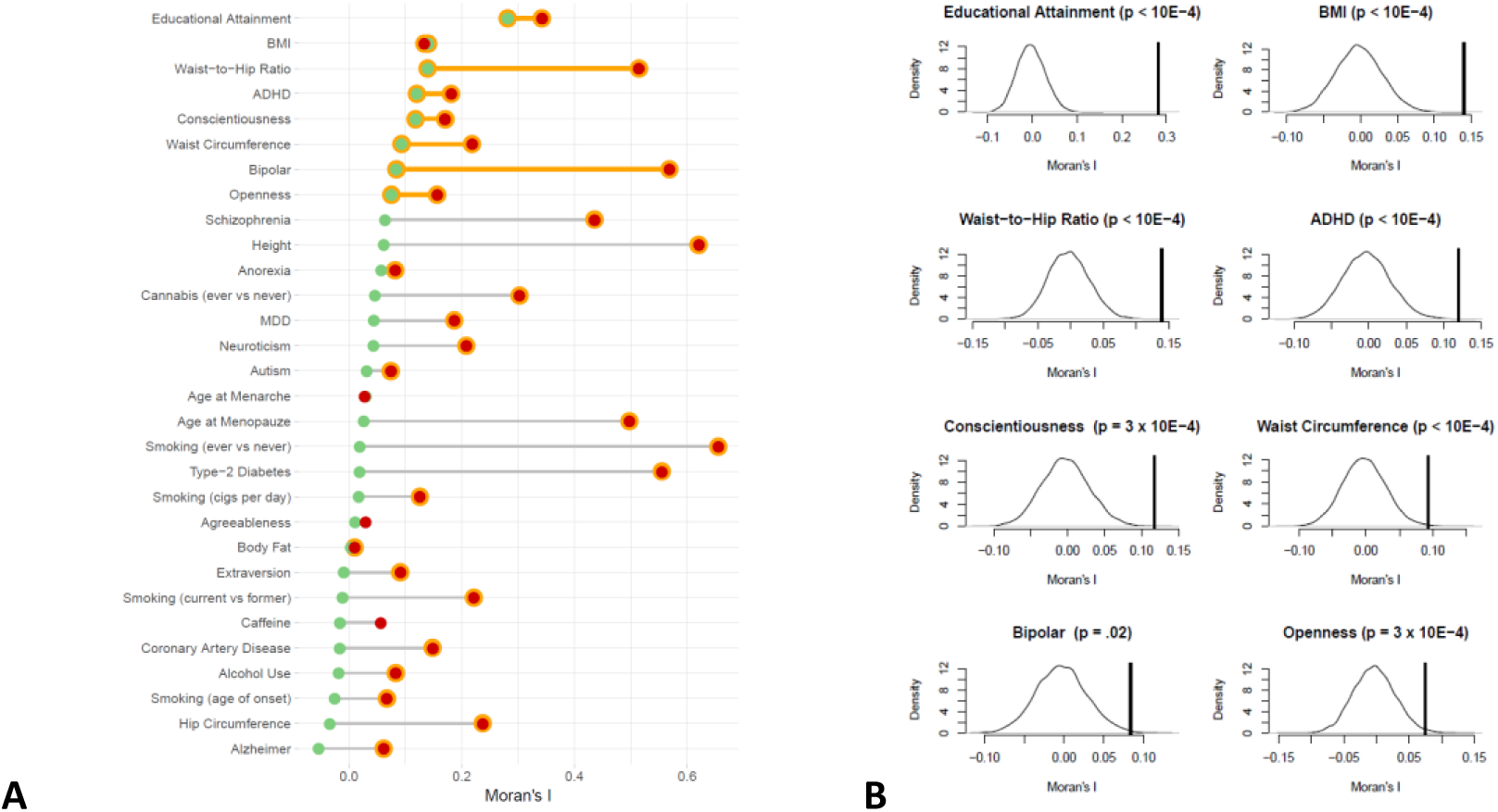
Moran’s *I* of 30 polygenic scores based on independent SNPs with *p* < .05. Moran’s *I* were computed using the average polygenic score per region in 378 local authority regions. **A** shows the Moran’s *I* of the polygenic scores uncorrected for PCs (red) and corrected for 100 PCs (green), where orange means a significant FDR corrected *p*-value < .05 (corrected for 30 tests). **B** shows the permutation distributions for the SBLUP polygenic scores that have an FDR corrected *p*-value < .05 compared to the observed Moran’s *I* (vertical line to the right of the permutation distribution).

**Supplementary Figure 3:**
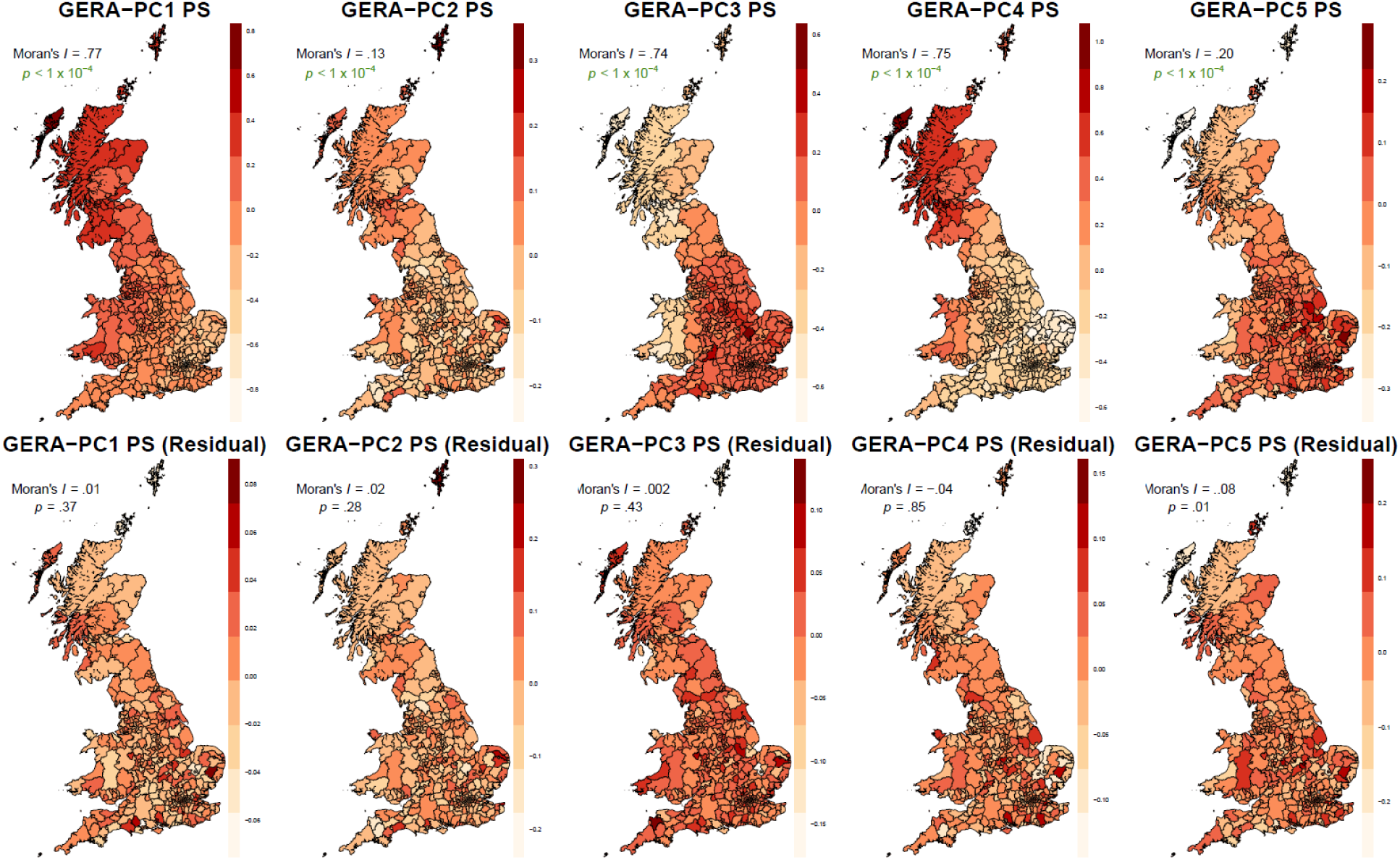
Geographic distribution and Moran’s *I* values for polygenic scores (PS) based on GWASs on the first 5 ancestry-informative PCs from the GERA dataset. The upper five maps display uncorrected polygenic score, while the five maps below display the residuals of the polygenic scores after regressing out 100 PCs. Green *p*-values are significant after FDR correction.

**Supplementary Figure 4:**
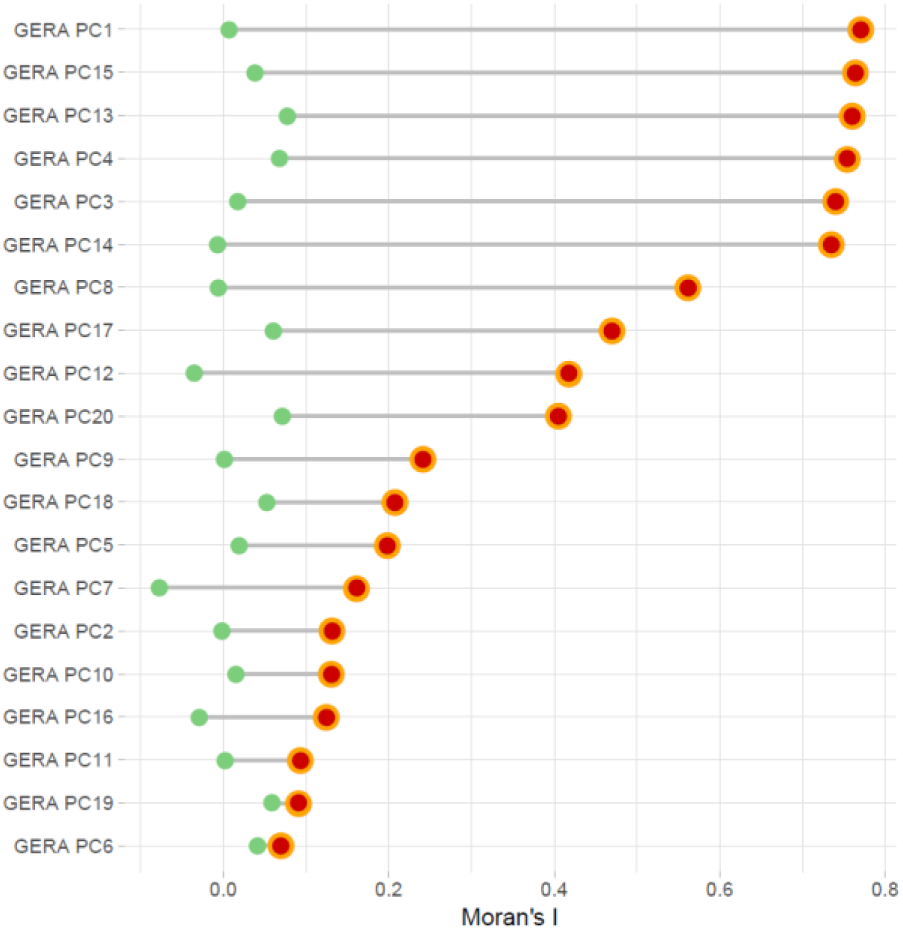
Moran’s *I* of polygenic scores based on GWASs on the first 5 ancestry-informative PCs from the GERA dataset computed using the average polygenic score per region in 378 local authority regions. The Figure shows the Moran’s *I* of the polygenic scores uncorrected for PCs (red) and corrected for 100 PCs (green), where orange means a significant FDR corrected *p*-value < .05 (corrected for 20 tests).

**Supplementary Figure 5:**
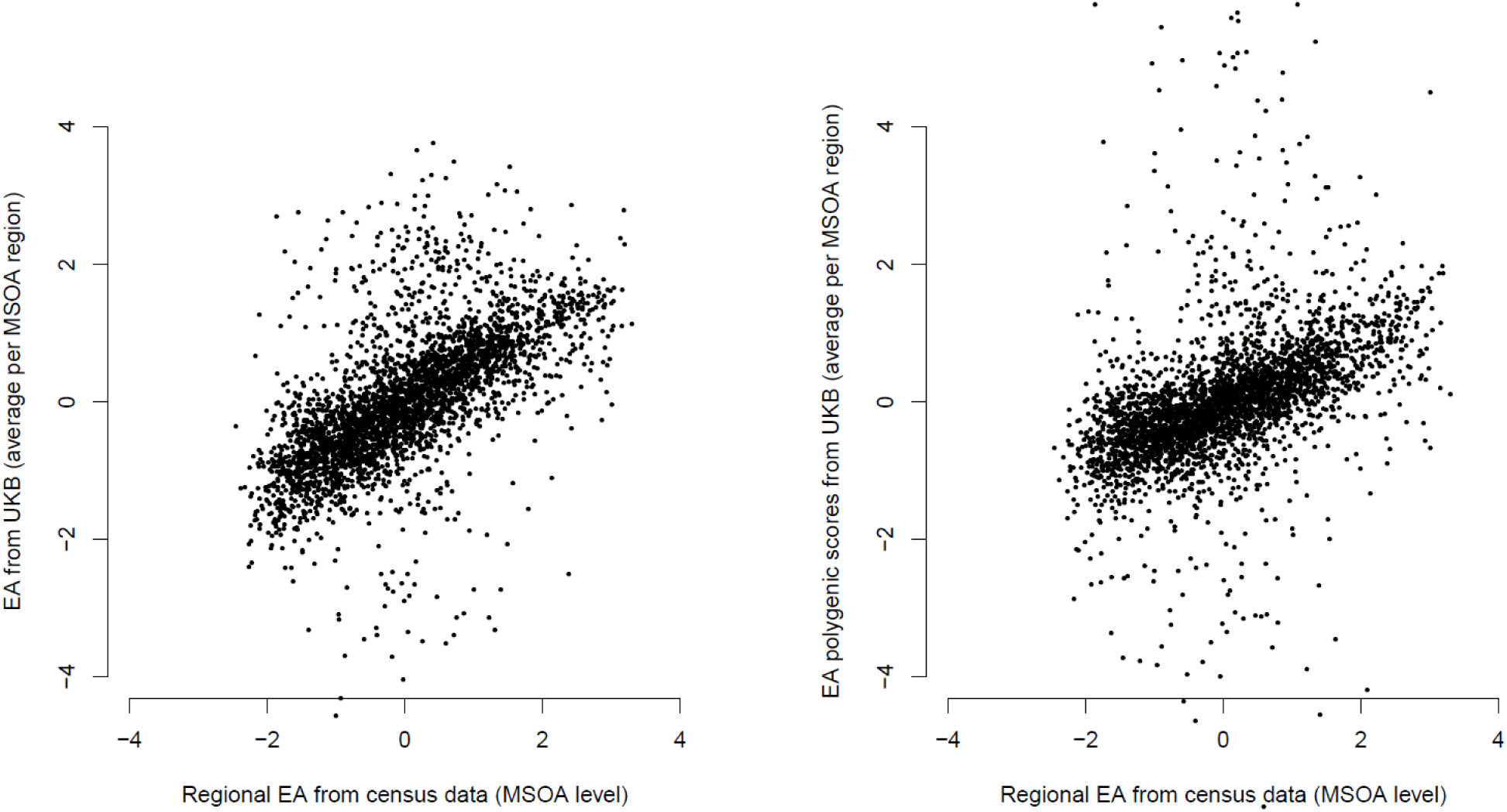
Regional educational attainment (EA) on MSOA level obtained from census data (Office of National Statistics) plotted against the average EA in UK Biobank corrected for age, year of birth, and sex (left plot; R^2^ = 40%) and EA polygenic scores corrected for 100 PCs (right plot; *R*^2^ = 20%). All measures have been standardized to have mean 0 and SD 1.

**Supplementary Figure 6:**
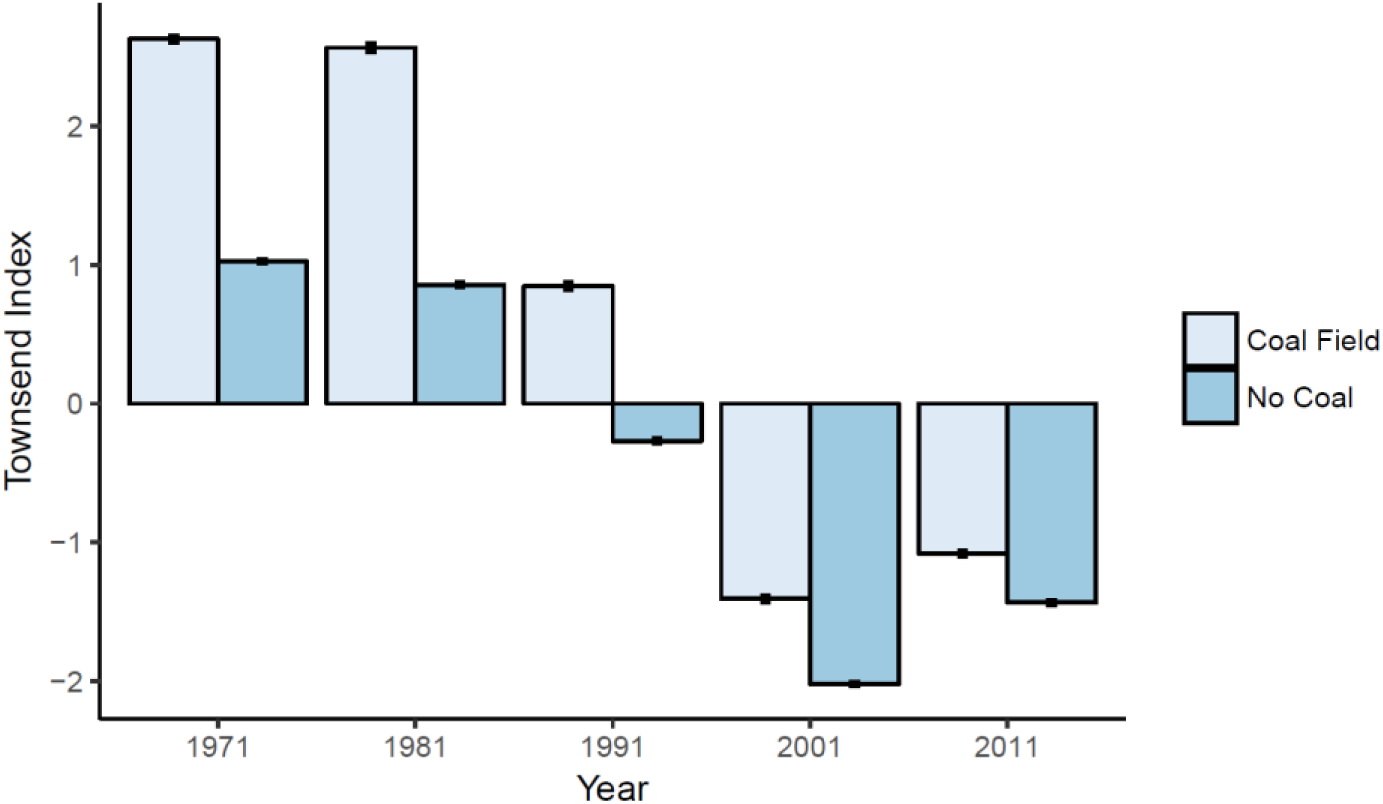
The average Townsend indices from 1971 to 2011 for coal fields and regions without coal.

**Supplementary Figure 7:**
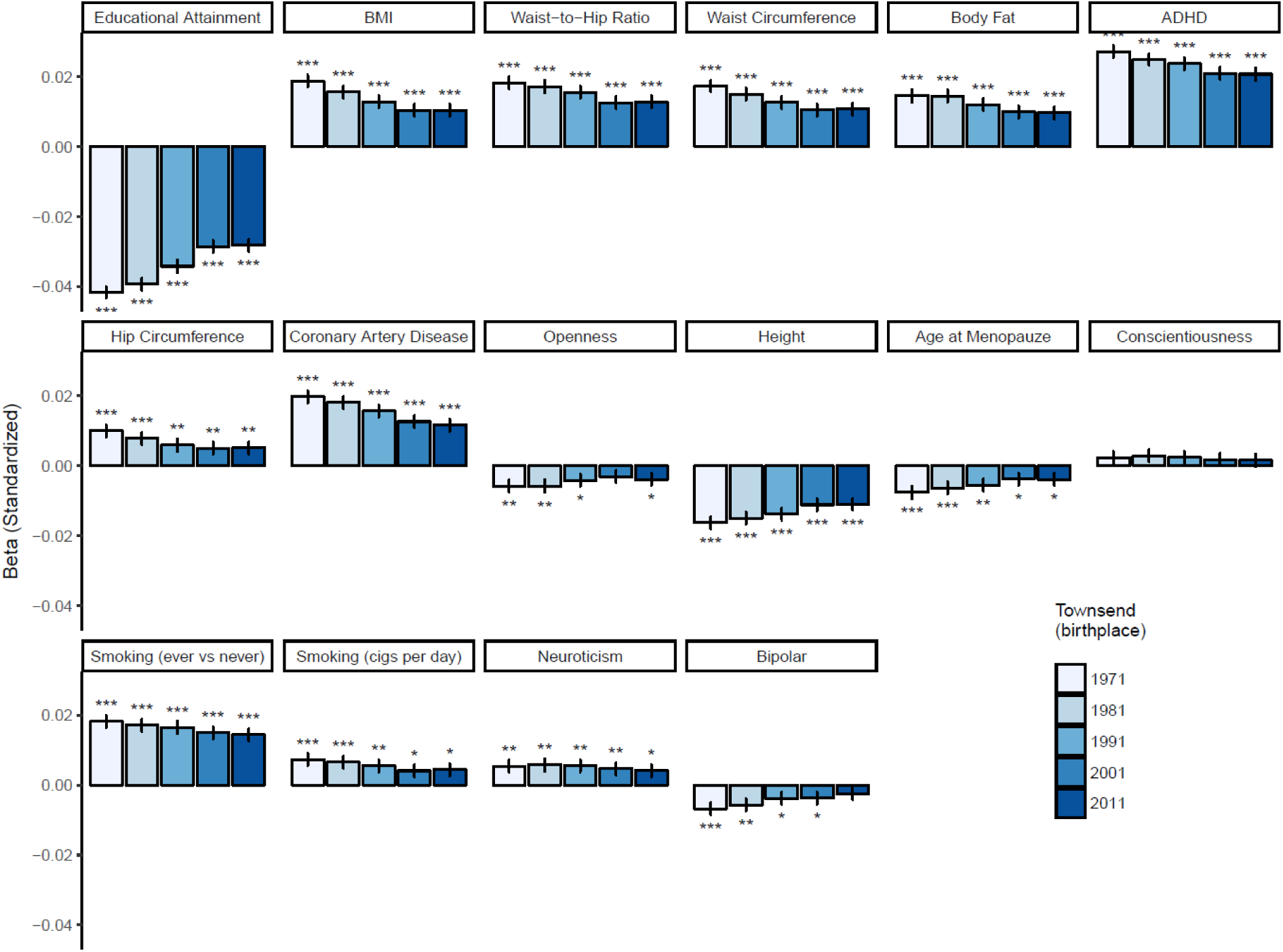
The standardized regression coefficients and standard errors for the 16 geographically clustered polygenic scores (ordered by Moran’s *I*) of the associations with the Townsend indices of 1971 to 2011 of the birth places of the subjects (N = 349,982 unrelated subjects). All polygenic scores shown are standardized residuals after regressing out 100 PCs. * = *p* < .05, ** = *p* < .01, *** = *p* < .001.

**Supplementary Figure 8:**
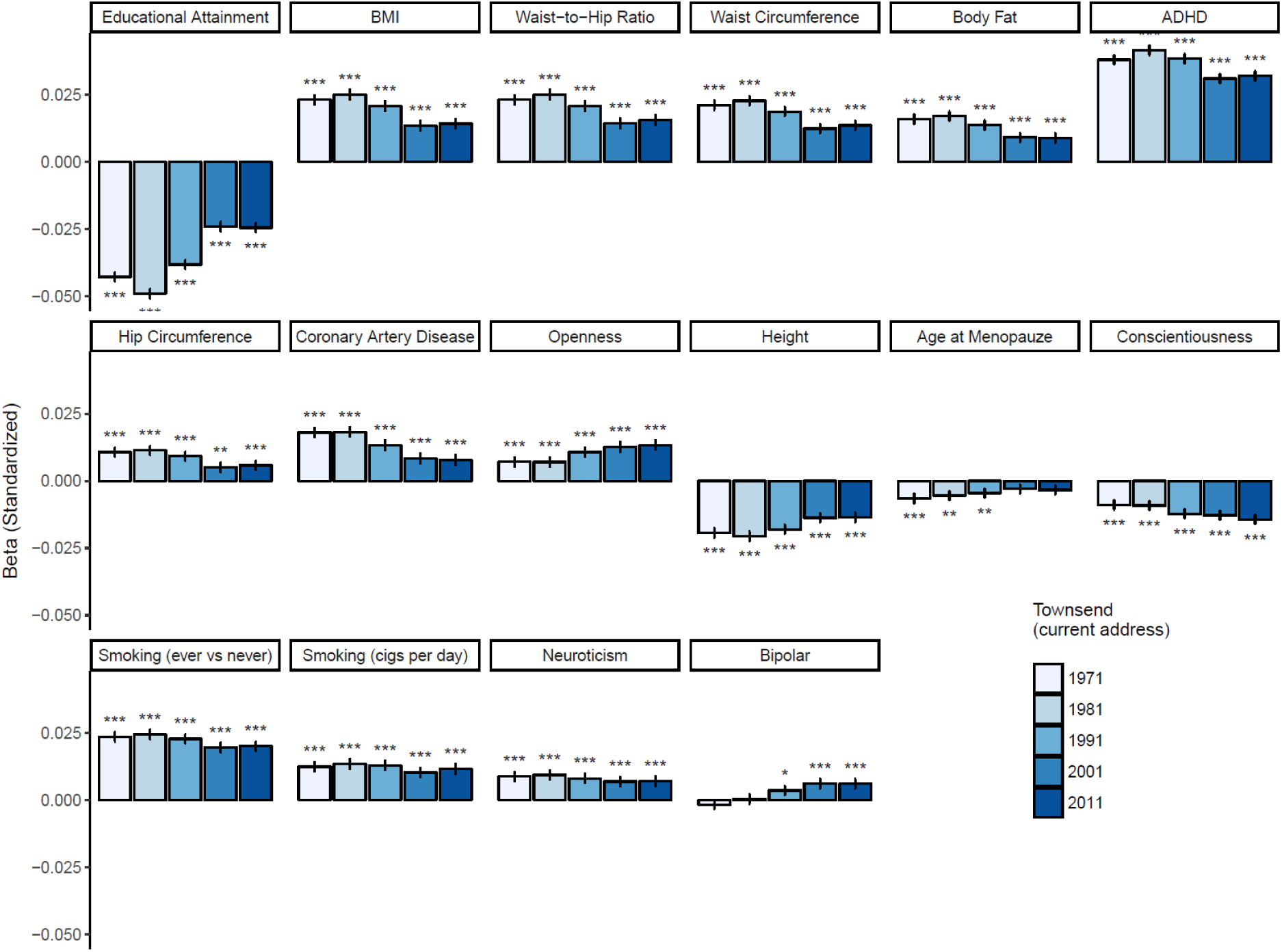
The standardized regression coefficients and standard errors for the 16 geographically clustered polygenic scores (ordered by Moran’s *I*) of the associations with the Townsend indices of 1971 to 2011 of the current address of the subjects (N = 349,982 unrelated subjects). All polygenic scores shown are standardized residuals after regressing out 100 PCs. * = *p* < .05, ** = *p* < .01, *** = *p* < .001.

**Supplementary Figure 9:**
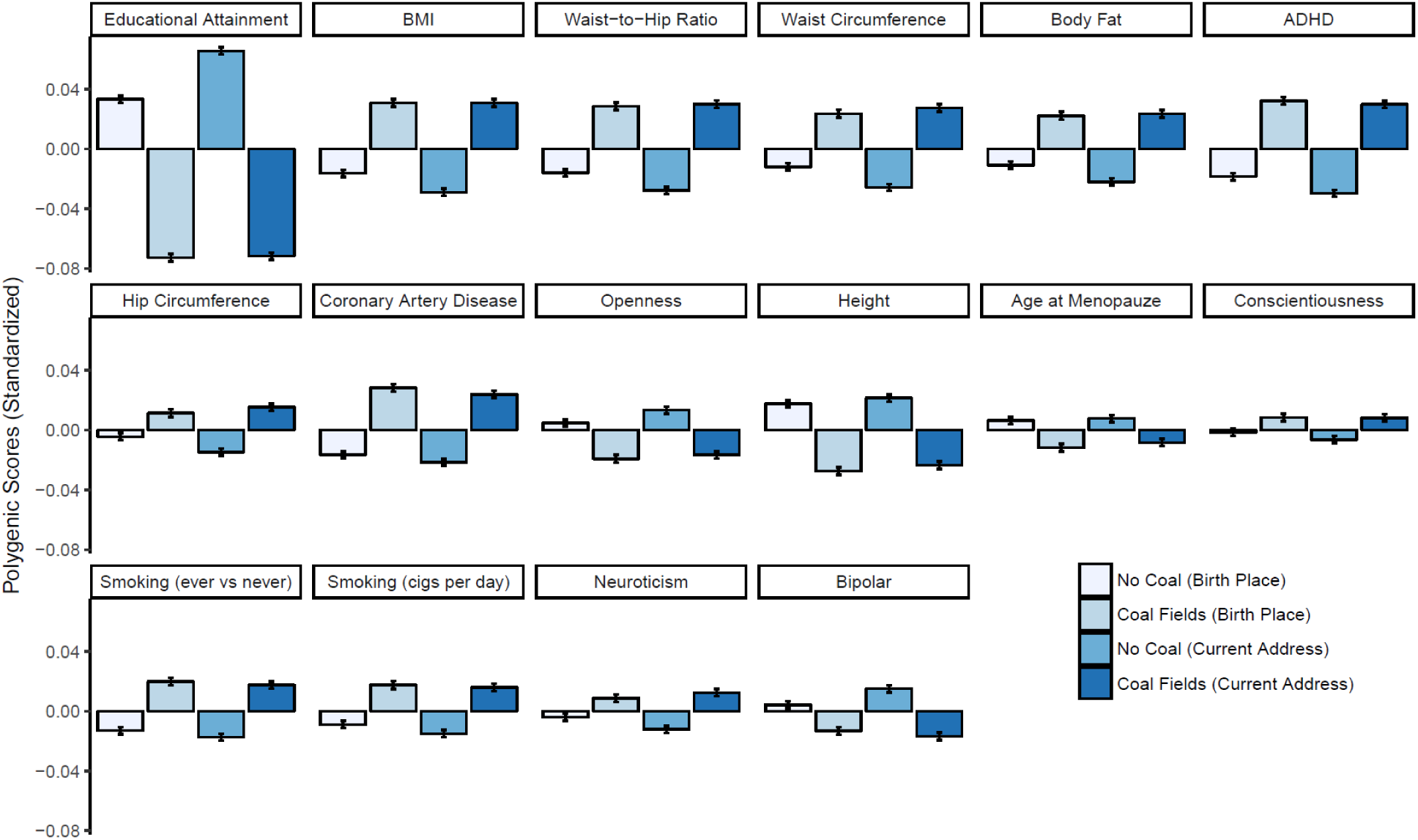
The average and standard errors for the 16 geographically clustered polygenic scores (ordered by Moran’s *I*) within coal mining regions vs the rest of Great Britain based on the birth place of the participants and on the current addresses separately. All polygenic scores shown are standardized residuals after regressing out 100 ancestry-informative PCs. The differences between Coal Fields and No Coal are all significant with an FDR corrected *p*-value < .05.

**Supplementary Figure 10:**
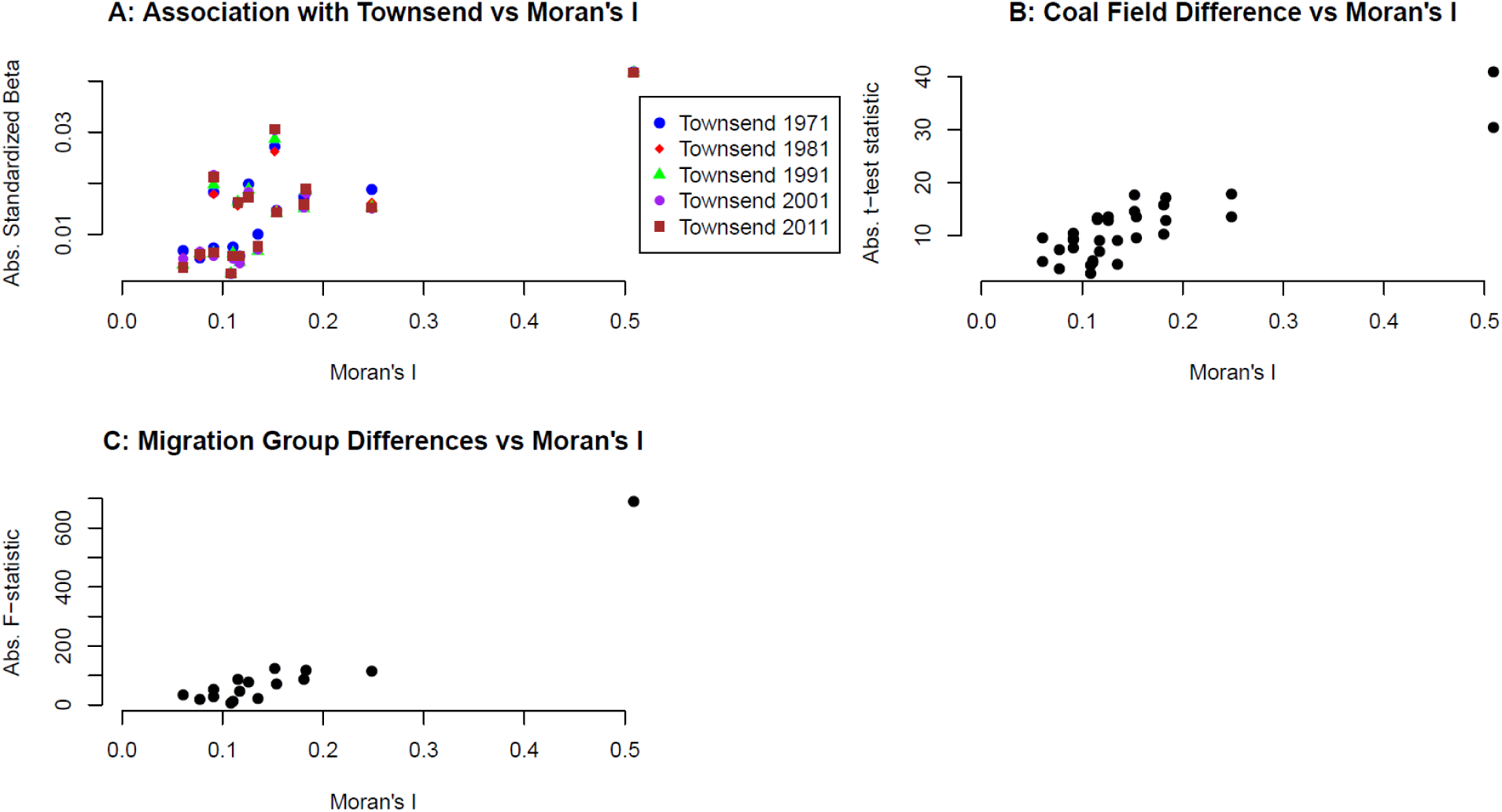
Scatterplots showing the relationships between Moran’s *I* of the significantly clustering polygenic scores and their association with indicators of economic deprivation (Townsend and the presence of coal fields) and migration. **A** shows on the *y*-axis the absolute regression coefficients of the regressions with birthplace Townsend shown in Supplementary Figure 3. The correlations between the absolute regression coefficients and Moran’s *I* are .82 for 1971 (*p* = 9 × 10^−5^), .82 for 1981 (*p* = 9 × 10^−5^), .78 for 1991 (*p* = 4 × 10^−4^), .75 for 2001 (*p* = 7 × 10^−4^), .77 for 2011 (*p* = 5 × 10^−4^). Excluding the outlier educational attainment, the correlations between the absolute regression coefficients and Moran’s *I* are .57 for 1971 (*p* = .03), .52 for 1981 (*p* = .04), .47 for 1991 (*p* = .08), .42 for 2001 (*p* = .12), .46 for 2011 (*p* = .08). **B** shows on the *y*-axis the absolute test statistic of the *t*-test for group differences between coal fields and the rest of Great Britain (based on birth place) from Supplementary Figure 5. The correlation between the absolute test statistic and Moran’s *I* is 89(*p* = 9 × 10^−12^). Excluding the outlier educational attainment, the correlations between the absolute test statistic and Moran’s *I* is .65 (*p* = 1 × 10^−4^). **C** shows on the *y*-axis the absolute test statistic of the ANOVA for group differences between the four migration groups from Figure 5. The correlation between the absolute test statistic and Moran’s *I* is .95 (*p* = 2 × 10^−8^). Excluding the outlier educational attainment, the correlations between the absolute test statistic and Moran’s *I* is .73 (*p* = .002).

**Supplementary Figure 11:**
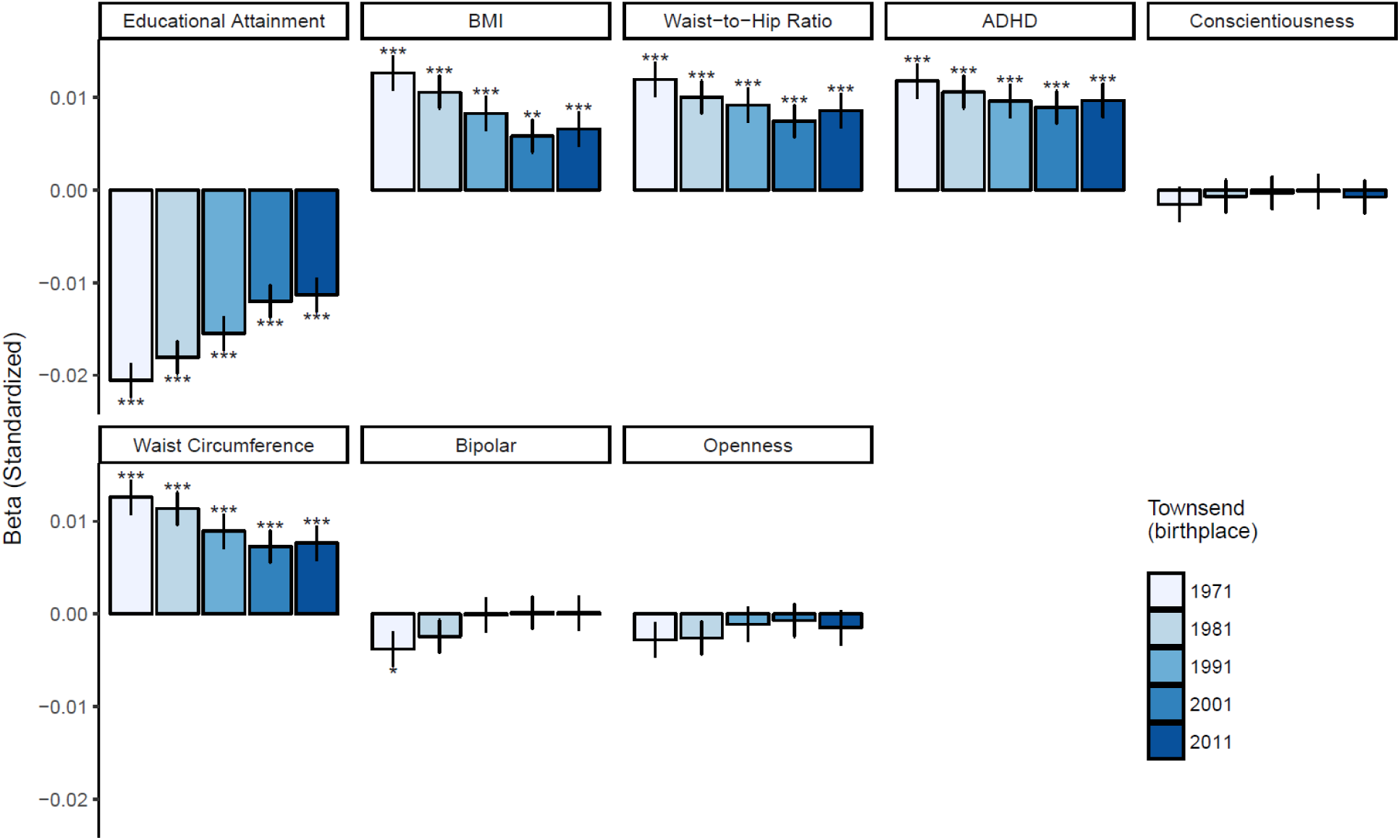
The standardized regression coefficients and standard errors for the 8 significantly clustering polygenic scores (clumped, i.e., based on independent SNPs with *p*-values <.05; ordered by Moran’s *I*) of the associations with the Townsend indices of 1971 to 2011 of the birth places of the subjects (N = 349,982 unrelated subjects). All polygenic scores shown are standardized residuals after regressing out 100 PCs.

**Supplementary Figure 12:**
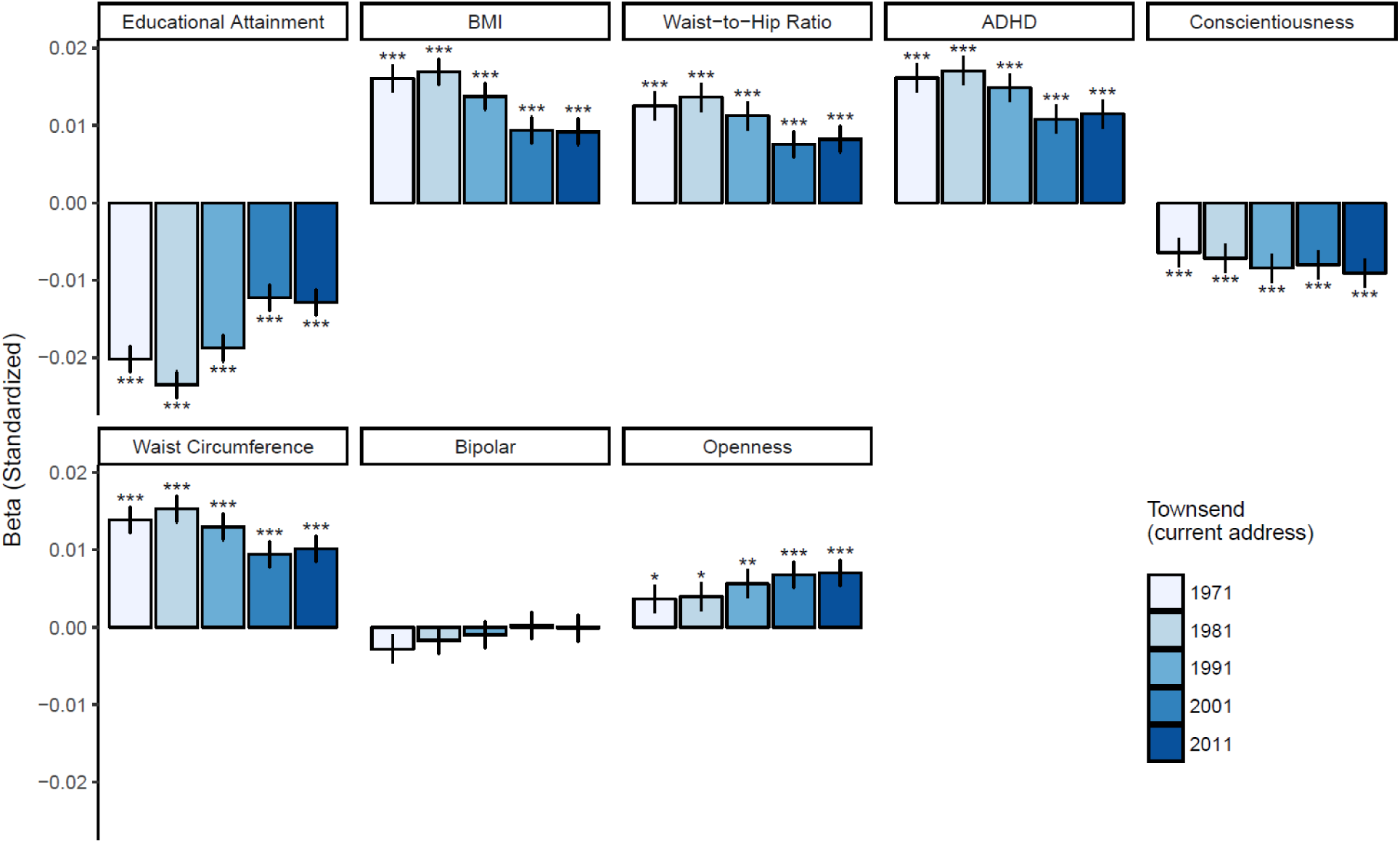
The standardized regression coefficients and standard errors for the 8 significantly clustering polygenic scores (clumped, i.e., based on independent SNPs with *p*-values <.05; ordered by Moran’s *I*) of the associations with the Townsend indices of 1971 to 2011 of the current address of the subjects (N = 349,982 unrelated subjects). All polygenic scores shown are standardized residuals after regressing out 100 PCs.

**Supplementary Figure 13:**
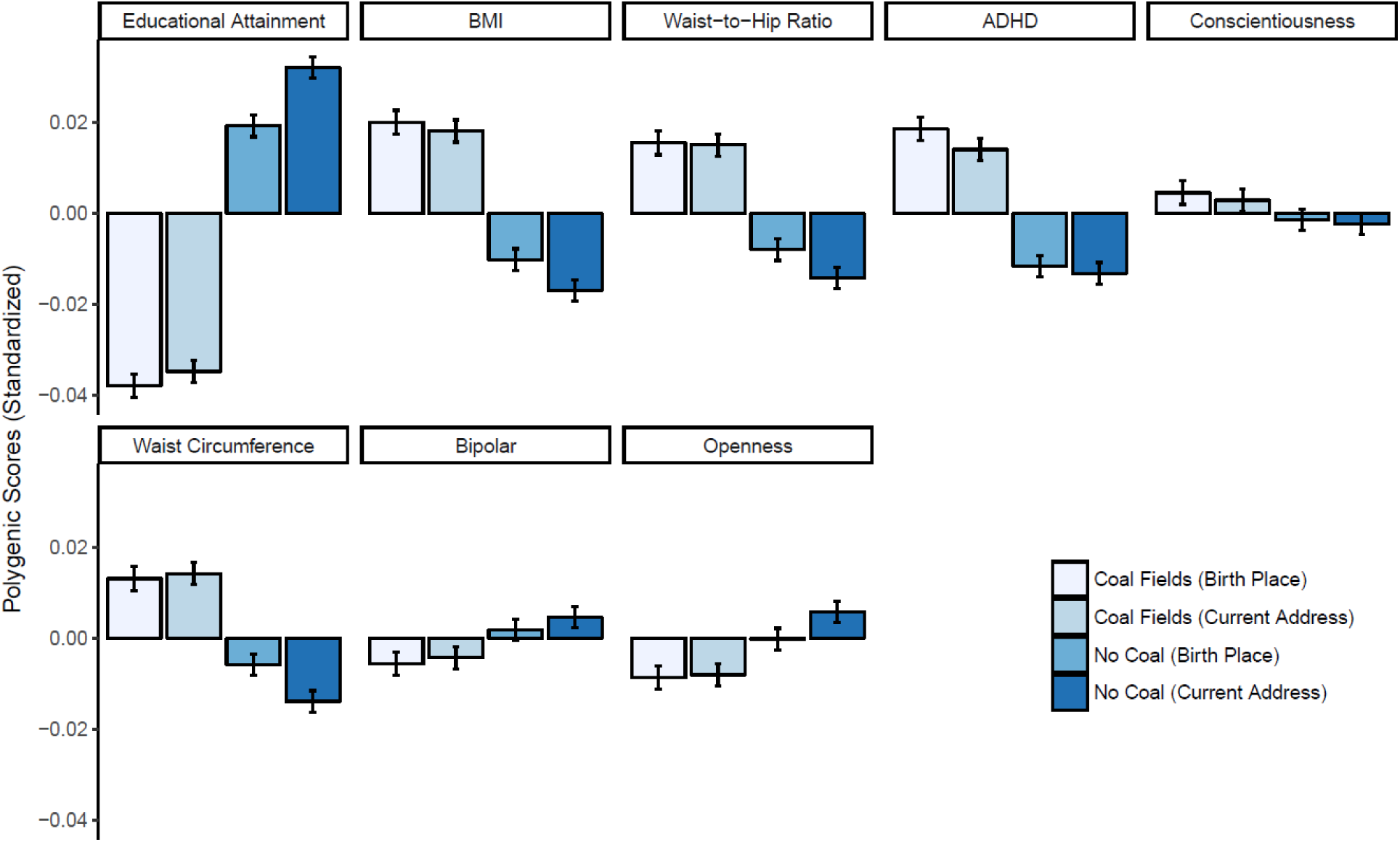
The average and standard errors for 8 significantly clustering polygenic scores (clumped, i.e., based on independent SNPs with *p*-values <.05; ordered by Moran’s *I*) within coal mining regions vs the rest of Great Britain for the birth place and the current address. All polygenic scores shown are standardized residuals after regressing out 100 ancestry-informative PCs. The differences between Coal Fields and No Coal are significant for all scores except conscientiousness for both current address and birth place with an FDR corrected *p*-value < .05.

**Supplementary Figure 14:**
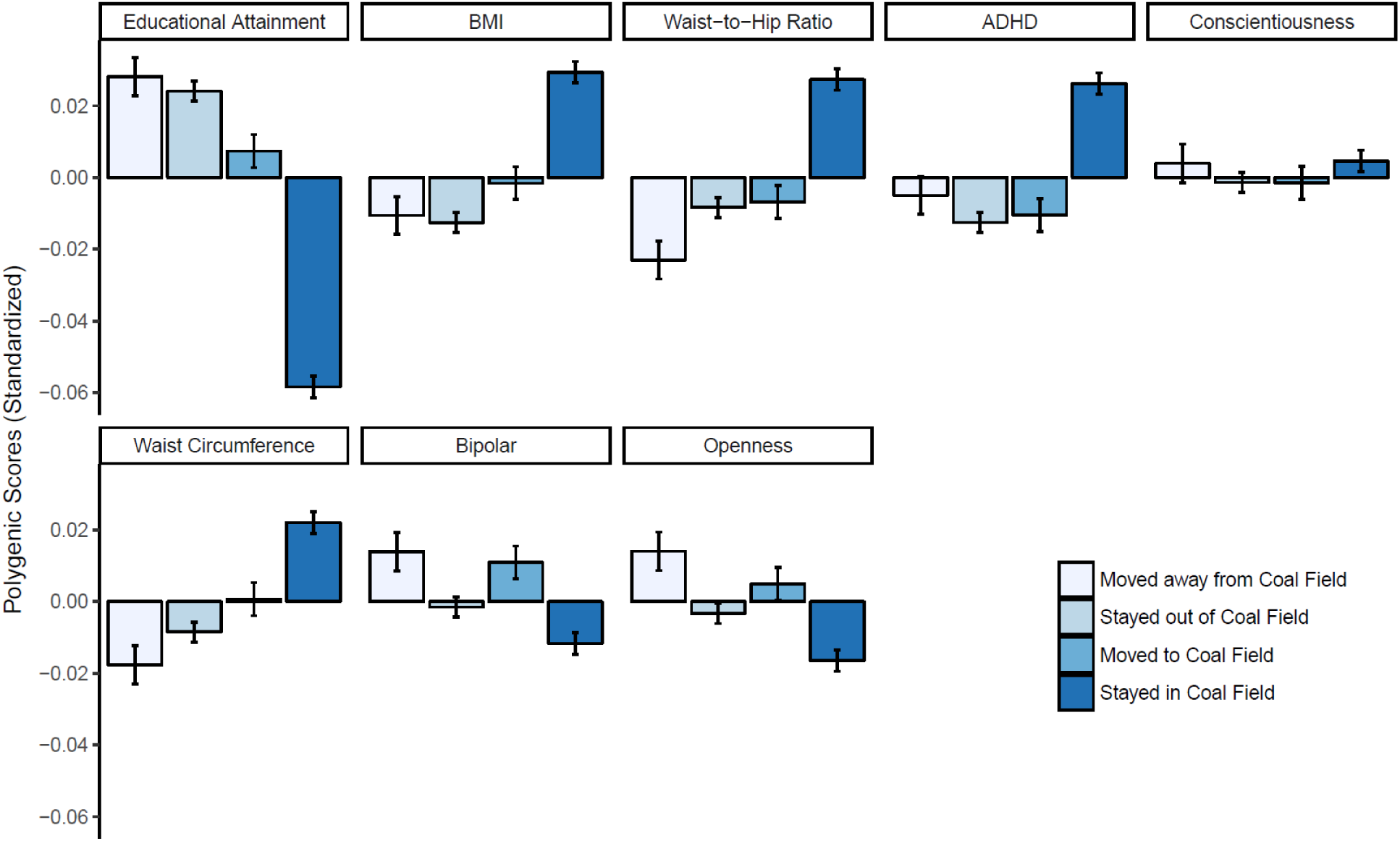
The average and standard errors for 8 significantly clustering polygenic scores (clumped, i.e., based on independent SNPs with *p*-values <.05; ordered by Moran’s *I*) for four migration groups: born in coal field area and moved out, born in coal field area and stayed, born outside of coal field area and moved to coal field area, born outside of coal field area and stayed out. All polygenic scores shown are standardized residuals after regressing out 100 ancestry-informative PCs. All polygenic scores show significant group differences after (FDR corrected *p* < .05), except conscientiousness.

**Supplementary Figure 15:**
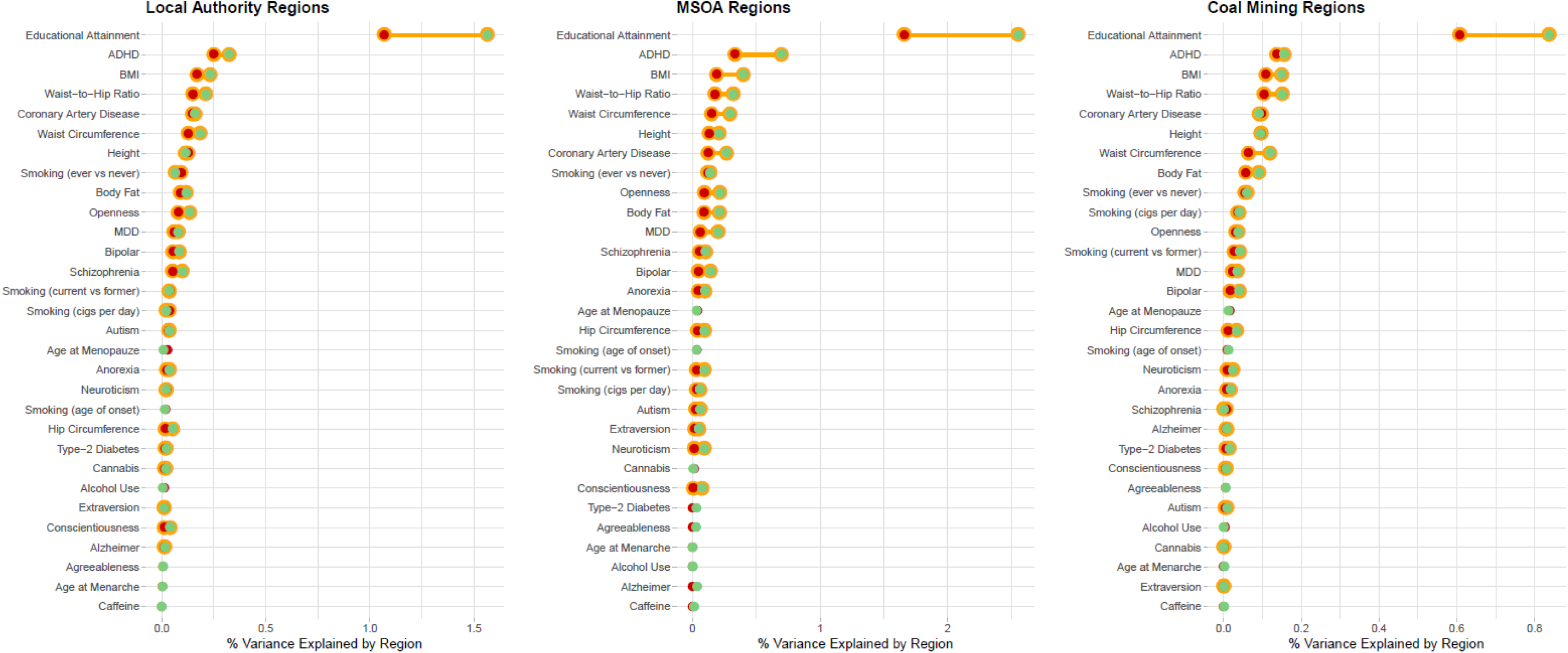
Linear Mixed Model results, with polygenic score (after regressing out 100 PCs) as a dependent variable and region as random effect (N = 320,940 unrelated individuals). Left: Local Authorities (~380 regions); Middle: MSOA (~5,300 regions), Right: Coal mining Regions (fitted as a binary variable). Red: Birth Place; Green: Current Address; Yellow = significant after FDR correction.

**Supplementary Figure 16:**
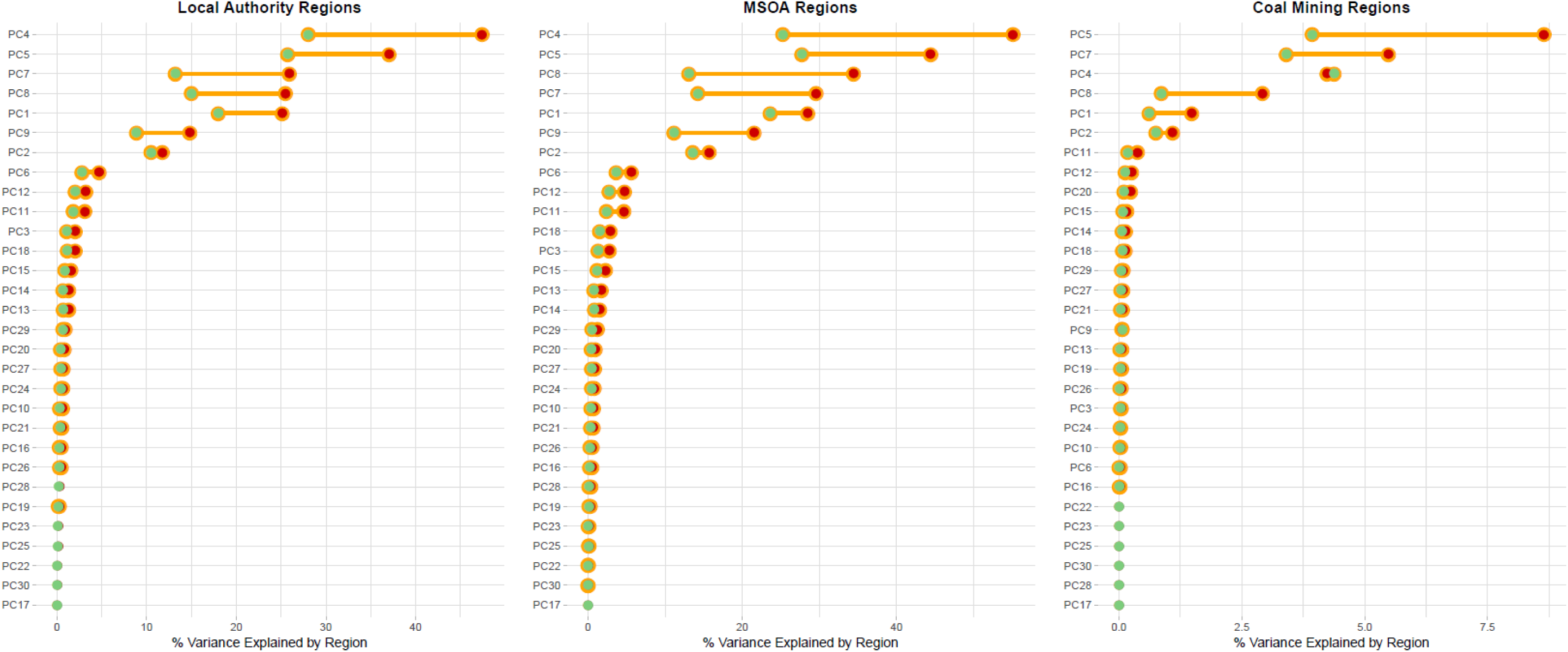
Linear Mixed Model results, with PCs as a dependent variable and region as random effect (N = 320,940 unrelated individuals). Left: Local Authorities (~380 regions); Middle: MSOA (~5,300 regions), Right: Coal mining Regions (fitted as a binary variable). Red: Birth Place; Green: Current Address; Yellow = significant after FDR correction.

**Supplementary Figure 17:**
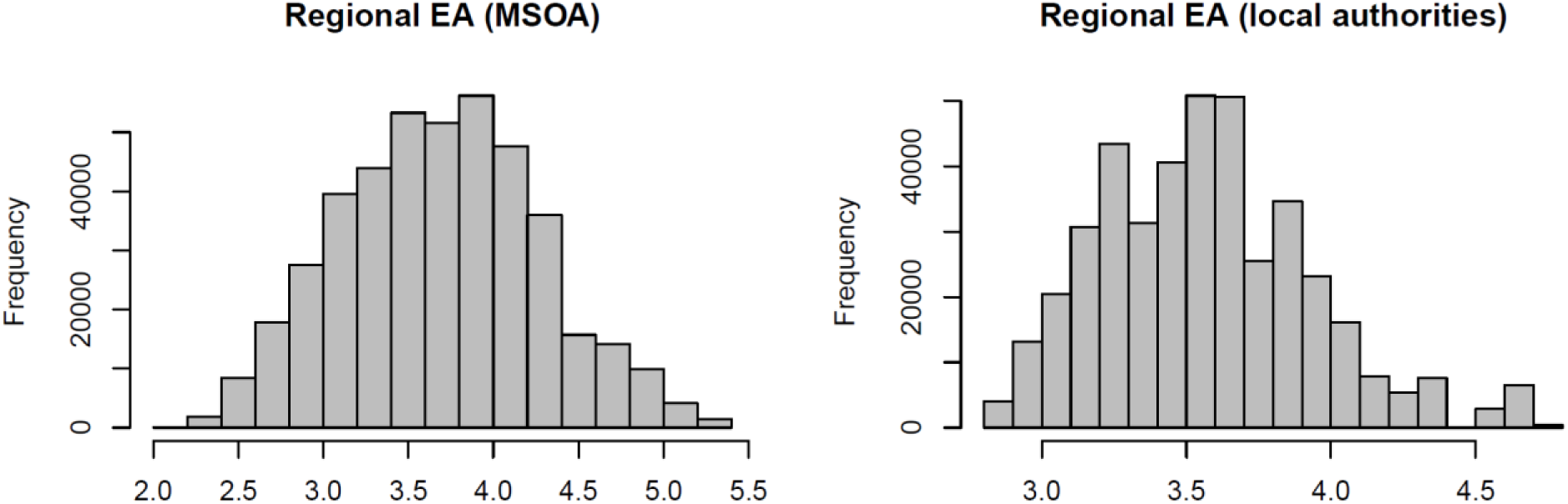
Distributions of the regional-level educational attainment (EA) phenotypes. The distributions show all subjects included in the GWASs, where all subjects from the same region were assigned the same phenotypic value.

**Supplementary Figure 18:**
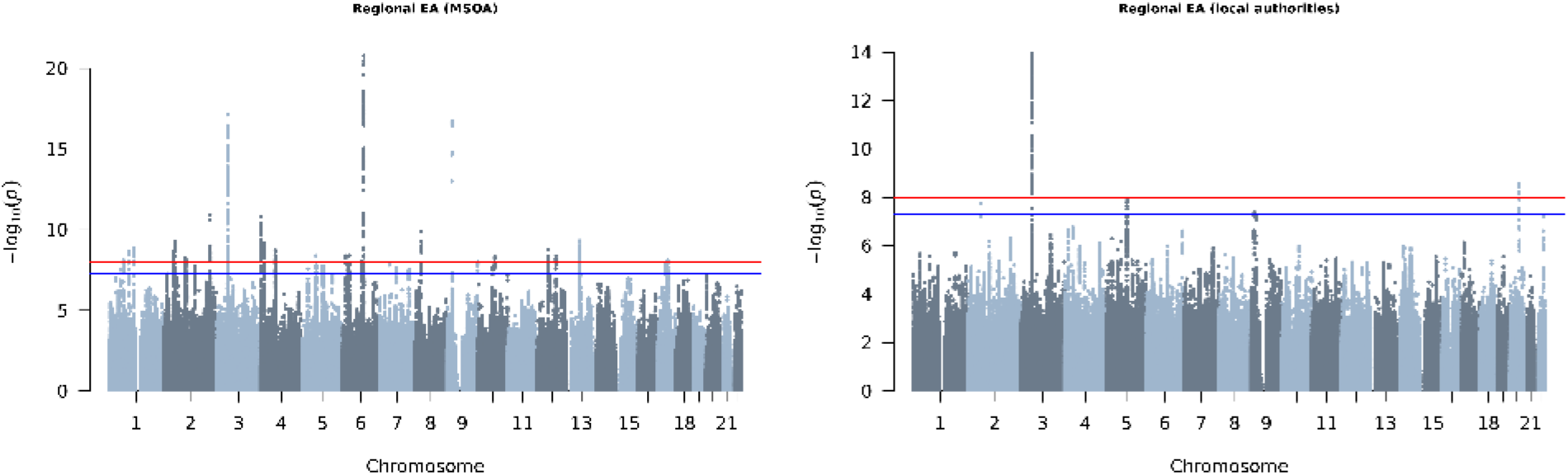
Manhattan plots of the two GC-corrected GWASs on regional educational attainment (EA). The suggestive significance threshold (blue line) is set at 5 × 10^−8^, and the genome-wide significance threshold (red line) is set at 1 × 10^−8^.

**Supplementary Figure 19:**
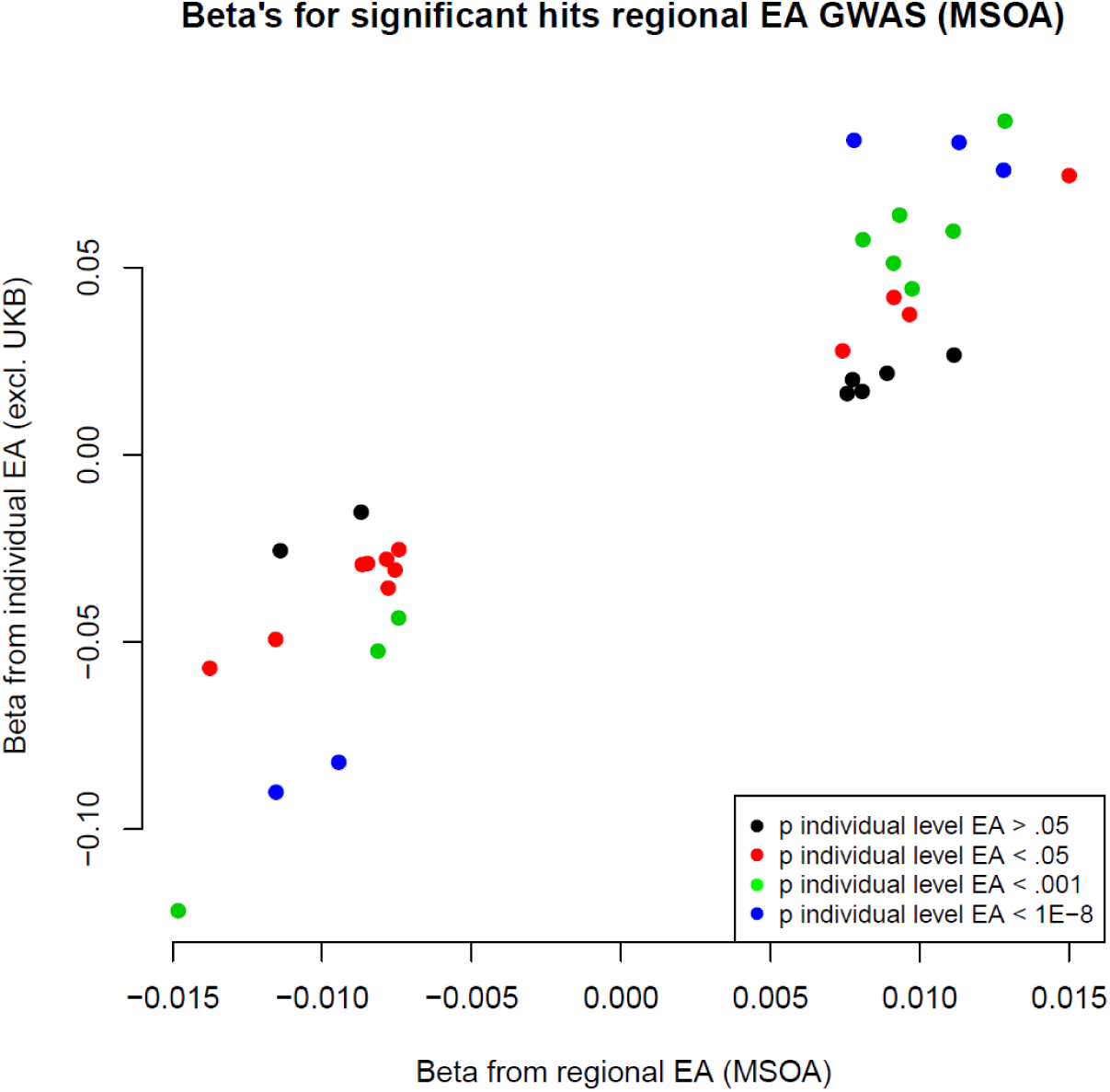
Effect sizes of 33 independent SNPs (*r*^2^ < .1 and at least 1MB apart) that reached *p* < 5 × 10^−8^ in the regional EA GWAS (MSOA) plotted against their effect sizes from the GWAS on individual EA that excluded UK Biobank participants. The correlation between the beta’s is .93.

**Supplementary Figure 20:**
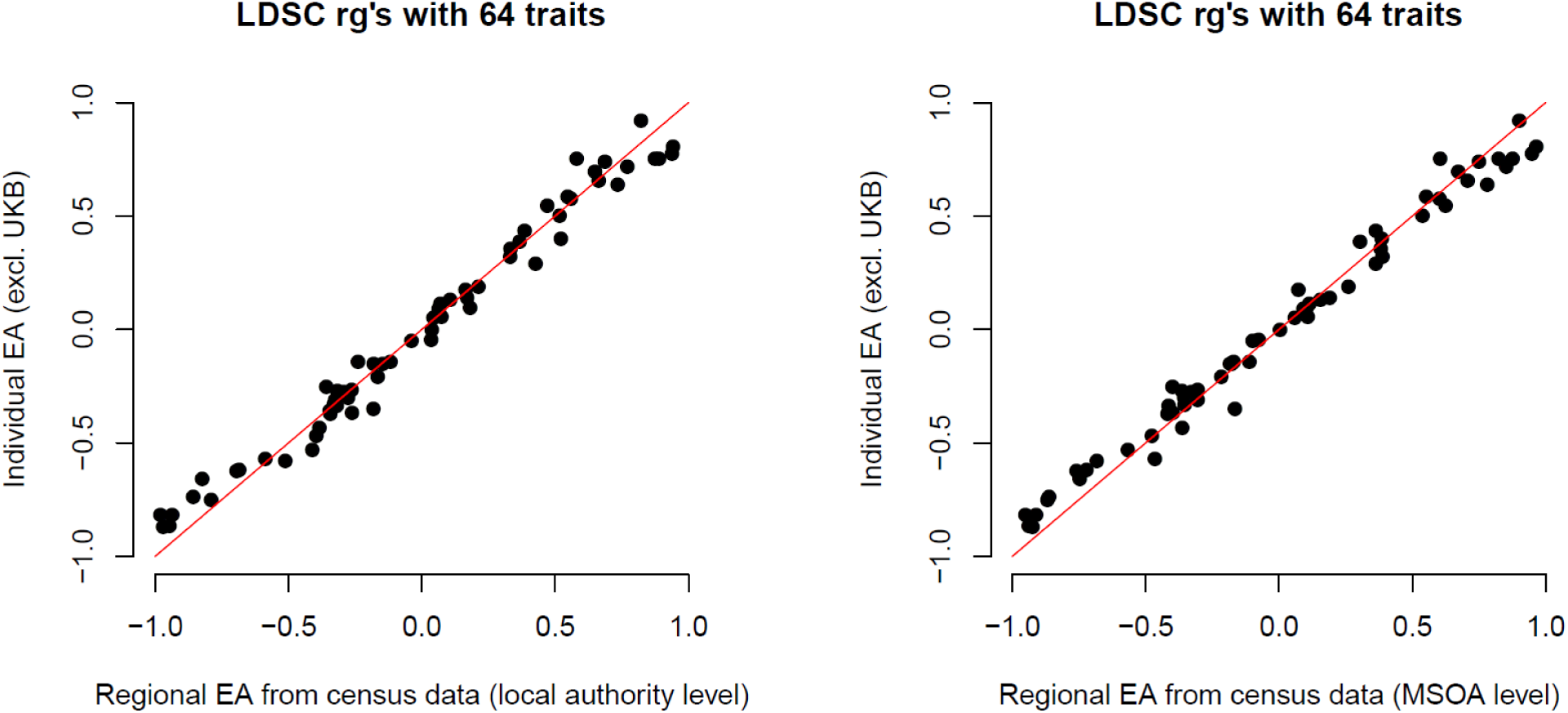
Scatterplots of the genetic correlations with 64 complex traits from Supplementary Figure 16. Each dot represents a complex trait. The correlations between the rg’s from the regional EA GWAS at local authority level and the individual EA GWAS (plot to the left) is .99. The correlations between the rg’s from the regional EA GWAS at MSOA level and the individual EA GWAS (plot to the right) is also .99.

**Supplementary Figure 21:**
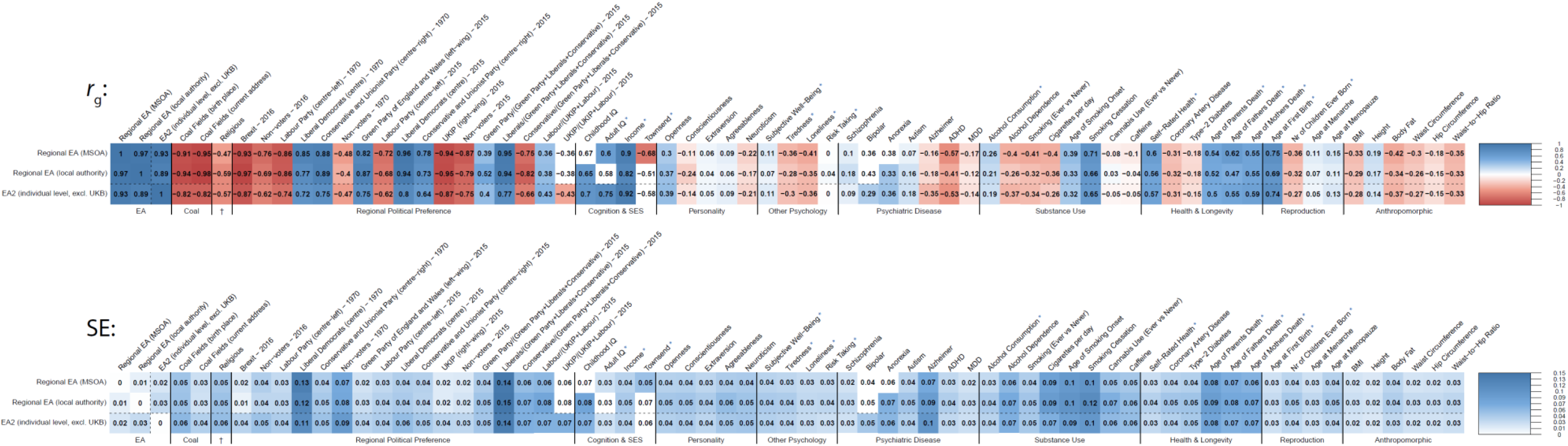
Genetic correlations (*r_g_*) and their standard errors (SE) based on LD score regression, for regional EA outcomes based on census data and the individual-level EA GWAS excluding UK Biobank. Colored is significant after FDR correction. The blue stars next to the trait names indicate that UK Biobank was part of the GWAS of the trait.

**Supplementary Figure 22:**
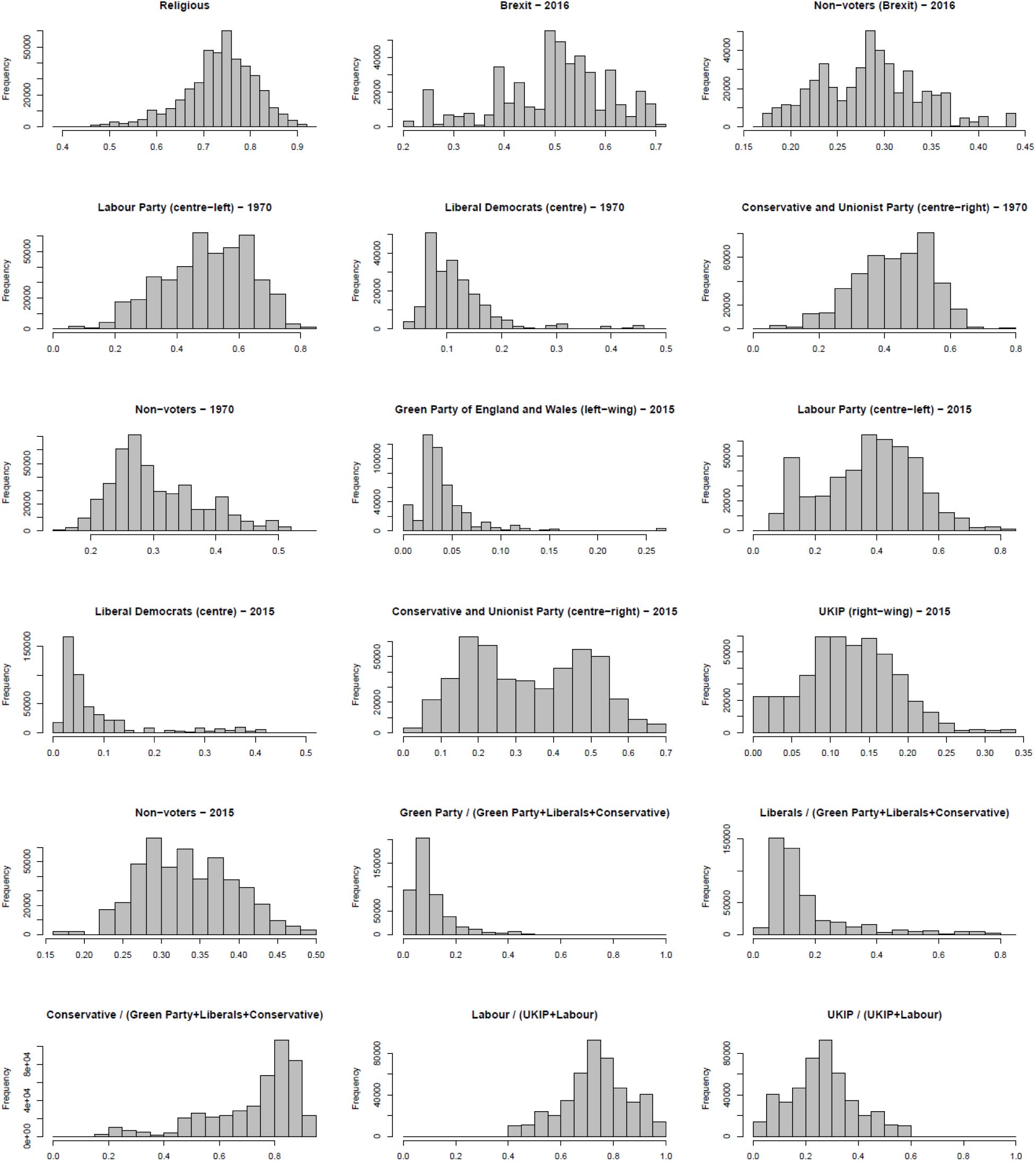
Distributions of regional-level phenotypes. The distributions show all subjects included in the GWASs, where all subjects from the same region were assigned the same phenotypic value.

**Supplementary Figure 23:**
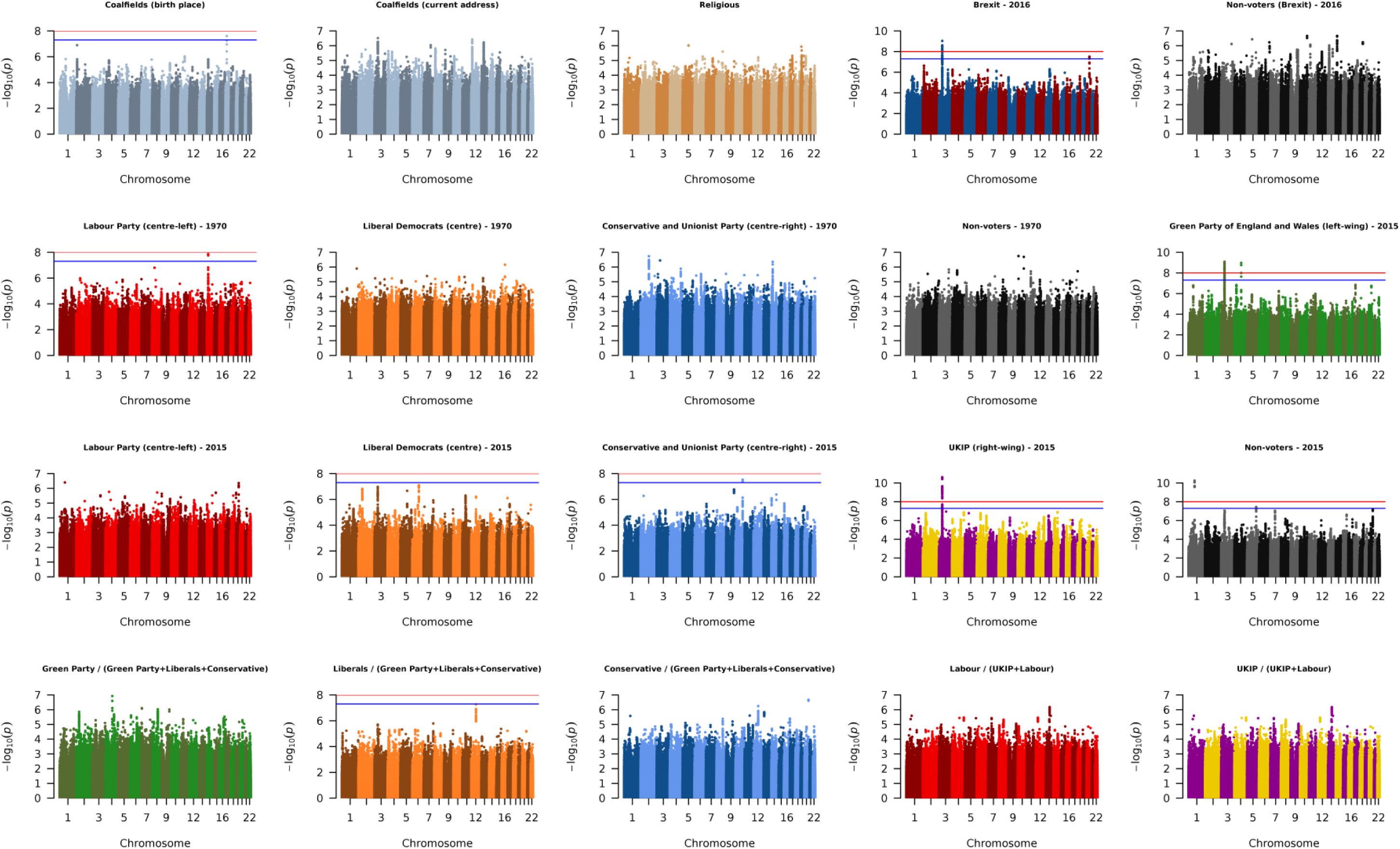
Manhattan plots of the GC-corrected GWAS on the presence of coal fields in the birthplace and the current address, the proportion of religious vs non-religious inhabitants, the Brexit referendum of 2016, and the regional outcomes on general election results of 1970 and 2015. The suggestive significance threshold (blue line) is set at 5 × 10^−8^, and the genome-wide significance threshold (red line) is set at 1 × 10^−8^.

**Supplementary Figure 24:**
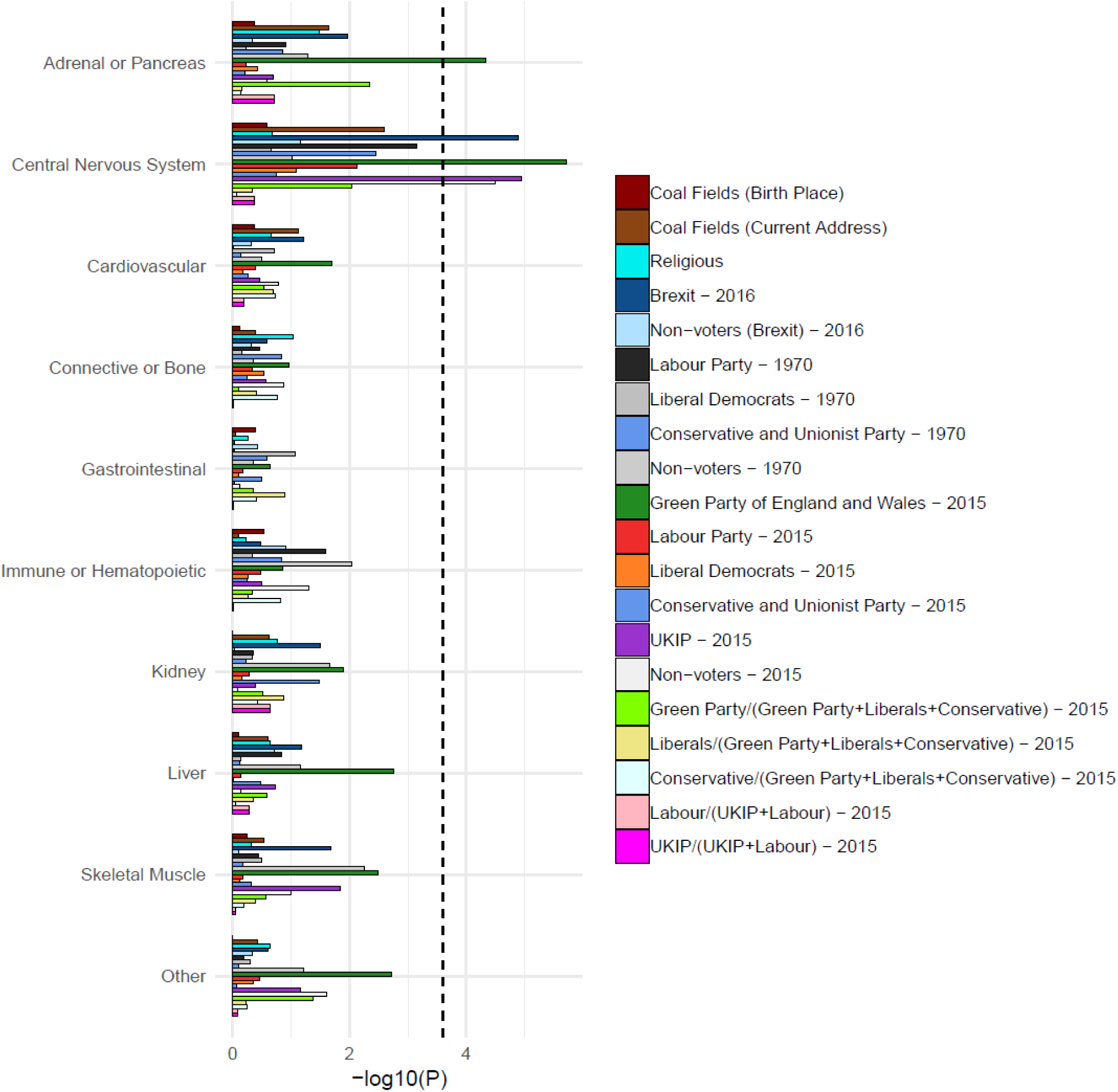
The significance of enrichment for ten cell type groups based on heritability partitioning with LD score regression. The dashed line at −*log*10(*P*) = 3.53 is the Bonferroni adjusted significance threshold (adjusted for 170 tests).

**Supplementary Figure 25:**
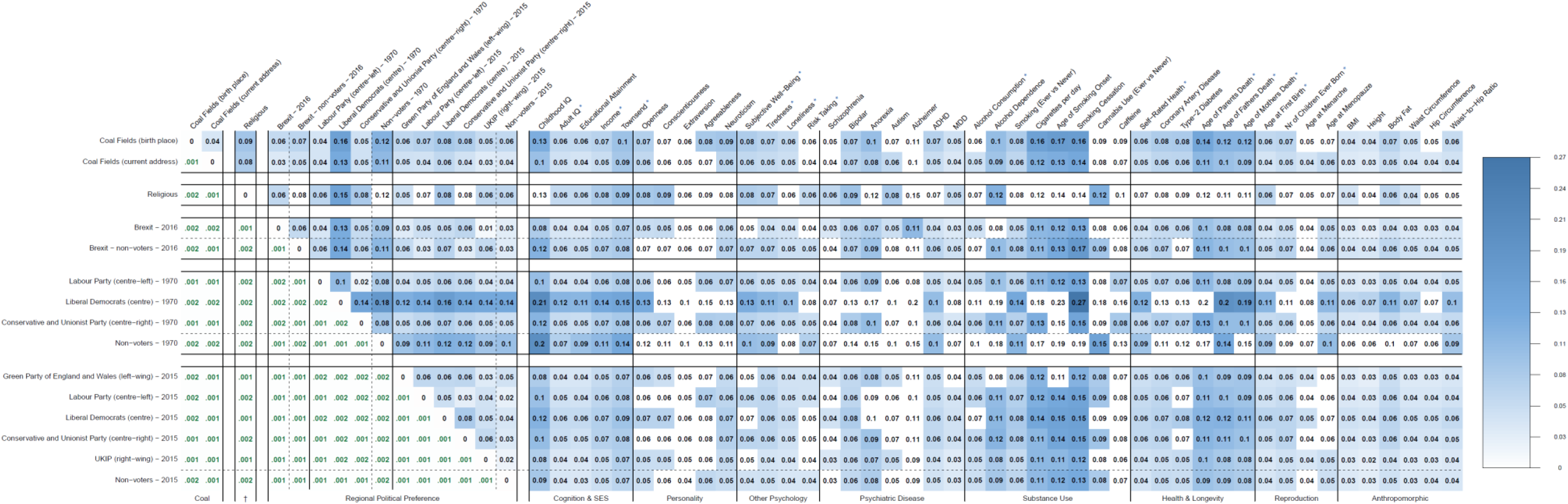
Standard errors of the genetic correlations based on LD score regression from Figure 6. Colored is significant after FDR correction. The green numbers in the left part of the Figure below the diagonal of 1’s are the standard errors of the phenotypic correlations between the regional outcomes of coal mining, religiousness, and regional political preference. The blue stars next to the trait names indicate that UK Biobank was part of the GWAS of the trait.

**Supplementary Figure 26:**
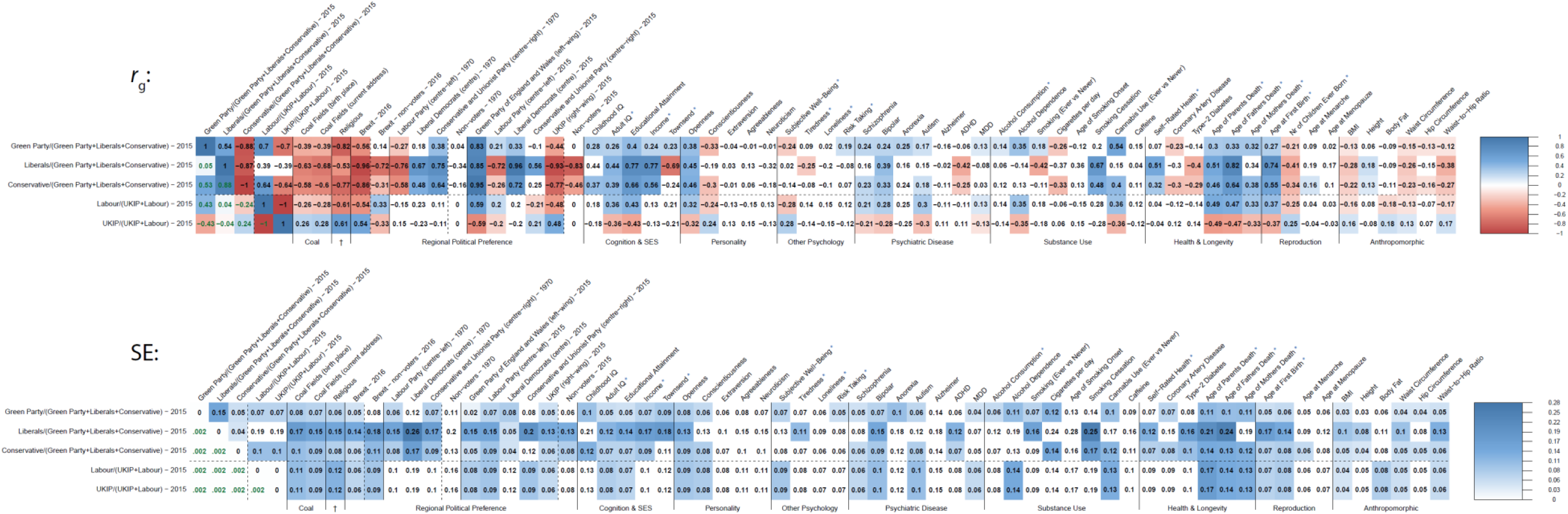
Genetic correlations (*r_g_*) and their standard errors (SE) based on LD score regression between regional political outcomes, within higher and lower SES groups. Colored is significant after FDR correction. The blue stars next to the trait names indicate that UK Biobank was part of the GWAS of the trait.

